# Estrogen therapy induces receptor-dependent DNA damage enhanced by PARP inhibition in ER+ breast cancer

**DOI:** 10.1101/2023.03.16.532956

**Authors:** Nicole A. Traphagen, Gary N. Schwartz, Steven Tau, Amanda Jiang, Sarah R. Hosford, Abigail E. Goen, Alyssa M. Roberts, Bianca A. Romo, Anneka L. Johnson, Emily-Claire K. Duffy, Eugene Demidenko, Paul Heverly, Yaron Mosesson, Shannon M. Soucy, Fred Kolling, Todd W. Miller

**Author notes:** **Corresponding Author** Todd W. Miller, Dartmouth-Hitchcock Medical Center, One Medical Center Drive, HB-7936, Lebanon, NH 03756. **Statement of Translational Relevance** A subset of patients with endocrine-resistant estrogen receptor alpha (ER)-positive breast cancer benefit from treatment with estrogens. However, the molecular effects and anti-cancer mechanism of action of estrogen therapy are unclear, which has limited the clinical use of this seemingly paradoxical treatment. We show that therapeutic response to the estrogen 17β-estradiol is dependent upon re-engagement of ER, and that 17β-estradiol treatment induces ER-dependent DNA damage in cells adapted to growth without estrogens. Pharmacological inhibition of poly(ADP-ribose) polymerase (PARP) synergizes with 17β-estradiol to enhance DNA-damage and therapeutic response. Importantly, this synergistic effect was observed regardless of BRCA1/2 mutation status. These findings collectively offer 17β-estradiol and PARP inhibitor combination treatment as a novel therapeutic strategy for patients with advanced ER+ breast cancer. Moreover, these data indicate that PARP inhibitors may have applications beyond homologous recombination-deficient tumors.

## Abstract

**Purpose:** Clinical evidence indicates that treatment with estrogens elicits anti-cancer effects in ∼30% of patients with advanced endocrine-resistant estrogen receptor alpha (ER)-positive breast cancer. Despite the proven efficacy of estrogen therapy, its mechanism of action is unclear and this treatment remains under-utilized. Mechanistic understanding may offer strategies to enhance therapeutic efficacy.

**Experimental Design:** We performed genome-wide CRISPR/Cas9 screening and transcriptomic profiling in long-term estrogen-deprived (LTED) ER+ breast cancer cells to identify pathways required for therapeutic response to the estrogen 17β-estradiol (E2). We validated findings in cell lines, patient-derived xenografts (PDXs), and patient samples, and developed a novel combination treatment through testing in cell lines and PDX models.

**Results:** Cells treated with E2 exhibited replication-dependent markers of DNA damage and the DNA damage response prior to apoptosis. Such DNA damage was partially driven by the formation of DNA:RNA hybrids (R-loops). Pharmacological suppression of the DNA damage response via poly(ADP-ribose) polymerase (PARP) inhibition with olaparib enhanced E2-induced DNA damage. PARP inhibition synergized with E2 to suppress growth and prevent tumor recurrence in *BRCA1/2*-mutant and *BRCA1*/2-wild-type cell line and PDX models.

**Conclusions:** E2-induced ER activity drives DNA damage and growth inhibition in endocrine-resistant breast cancer cells. Inhibition of the DNA damage response using drugs such as PARP inhibitors can enhance therapeutic response to E2. These findings warrant clinical exploration of the combination of E2 with DNA damage response inhibitors in advanced ER+ breast cancer, and suggest that PARP inhibitors may synergize with therapeutics that exacerbate transcriptional stress.

## Introduction

The majority of breast tumors express estrogen receptor α (ER), which typically reflects a degree of dependence upon estrogens for tumor growth. ER+/HER2-breast cancer is commonly treated with anti-estrogens that antagonize ER (e.g., tamoxifen, fulvestrant) or aromatase inhibitors that suppress estrogen biosynthesis. Although such endocrine therapies have improved outcomes for patients overall, endocrine resistance remains a clinical problem: approximately 20-30% of early-stage ER+ breast cancer patients experience disease recurrence. Despite the acquisition of resistance to anti-estrogens, loss of ER expression is rare in recurrent ER+ breast cancer and occurs in <10% of cases (1). Within the past 15 years, advances have been made in developing tumor-targeted therapies for endocrine-resistant disease (e.g., inhibitors of mTOR, CDK4/6, and PI3K). Since recurrent ER+ breast tumors remain at least partially dependent upon ER activity (2), approved tumor-targeted therapies are often administered in combination with an endocrine agent. Although these therapies have increased progression-free survival in patients, metastatic disease is typically fatal, and there remains a pressing need for new treatment options.

Decades of clinical evidence have demonstrated therapeutic efficacy of estrogen treatments in a subset of breast cancer patients (3-6). In the setting of endocrine-resistant advanced/metastatic ER+ disease, estrogens elicit anti-cancer effects in ∼30% of patients, translating into thousands of patients worldwide who could benefit from these treatments (7-11). Despite robust clinical evidence of efficacy, estrogen treatments remain under-utilized due in part to their unknown and seemingly paradoxical anti-cancer mechanism(s) of action. We previously demonstrated that therapeutic response to the natural estrogen 17β-estradiol (E2) requires ER, and hyperactivation of ER transcriptional activity through high levels of receptor expression and acute stimulation with ligand elicits anti-cancer effects (12). Herein, we demonstrate that estrogen therapy-induced apoptosis is dependent upon cell cycle progression. E2 induces ER-dependent DNA damage requiring R-loop formation and S-phase DNA replication, which can be exploited therapeutically through poly (ADP-ribose) polymerase (PARP) inhibition to enhance efficacy.

## Materials and Methods

### Cell Culture

Parental cell lines (ATCC) were cultured in DMEM with 10% FBS. For hormone deprivation (HD) experiments, cells were cultured in phenol red–free DMEM containing 10% dextran/charcoal-stripped FBS. Cells were stably transfected with lentiviral vectors encoding luciferase, FLAG-tagged ER (FLAG-*ESR1*), doxycycline (dox)-inducible ER (pInducer20-*ESR1*), dox-inducible shRNA targeting ER (*ESR1*), or non-targeting shControl (12). Cells were transiently transfected with plasmids encoding RNase H1 or vector control.

### Tumor growth studies

Animal studies were approved by the Dartmouth College IACUC. Female NOD-scid/IL2Rγ−/− (NSG) mice (4-6 wk) were ovariectomized and orthotopically implanted with fragments of WHIM16 or CTG-3346 patient-derived xenografts (PDX); WHIM16 was obtained from Washington University HAMLET Core (13). Tumor dimensions were measured twice weekly with calipers, and volumes were calculated as [length × width^2^/2]. When tumor volume reached ∼200 mm^3^, mice were randomized to treatments as indicated. For molecular analysis, tumors were harvested at the indicated time points and either snap frozen, or formalin-fixed and paraffin-embedded.

### Statistical analysis

Cell growth data and IHC scores were analyzed by t-test (for 2-group experiments) or ANOVA followed by Bonferroni multiple comparison-adjusted post hoc testing between groups (for experiments with ≥3 groups). Pairwise comparisons of immunofluorescence scores used the Cramer-von Mises nonparametric test. Tumor volumes were analyzed by nonlinear mixed modelling.

Additional details are provided in Supplemental Methods.

## Results

### Functional genomic screening suggests roles for cell cycle progression and DNA damage response in anti-cancer effects of 17β-estradiol

Parental ER+/HER2-HCC-1428 breast cancer cells are dependent upon E2 for growth (14). In contrast, their long-term estrogen-deprived (LTED) derivatives exhibit acquired resistance to hormone deprivation (HD) and growth is inhibited by re-treatment with 1 nM E2 [Fig. 1A and ref. (12)]. This concentration of E2 is within the pre-menopausal range in humans, and can be achieved pharmacologically in patients treated with E2 therapy (7).

**Figure 1.**
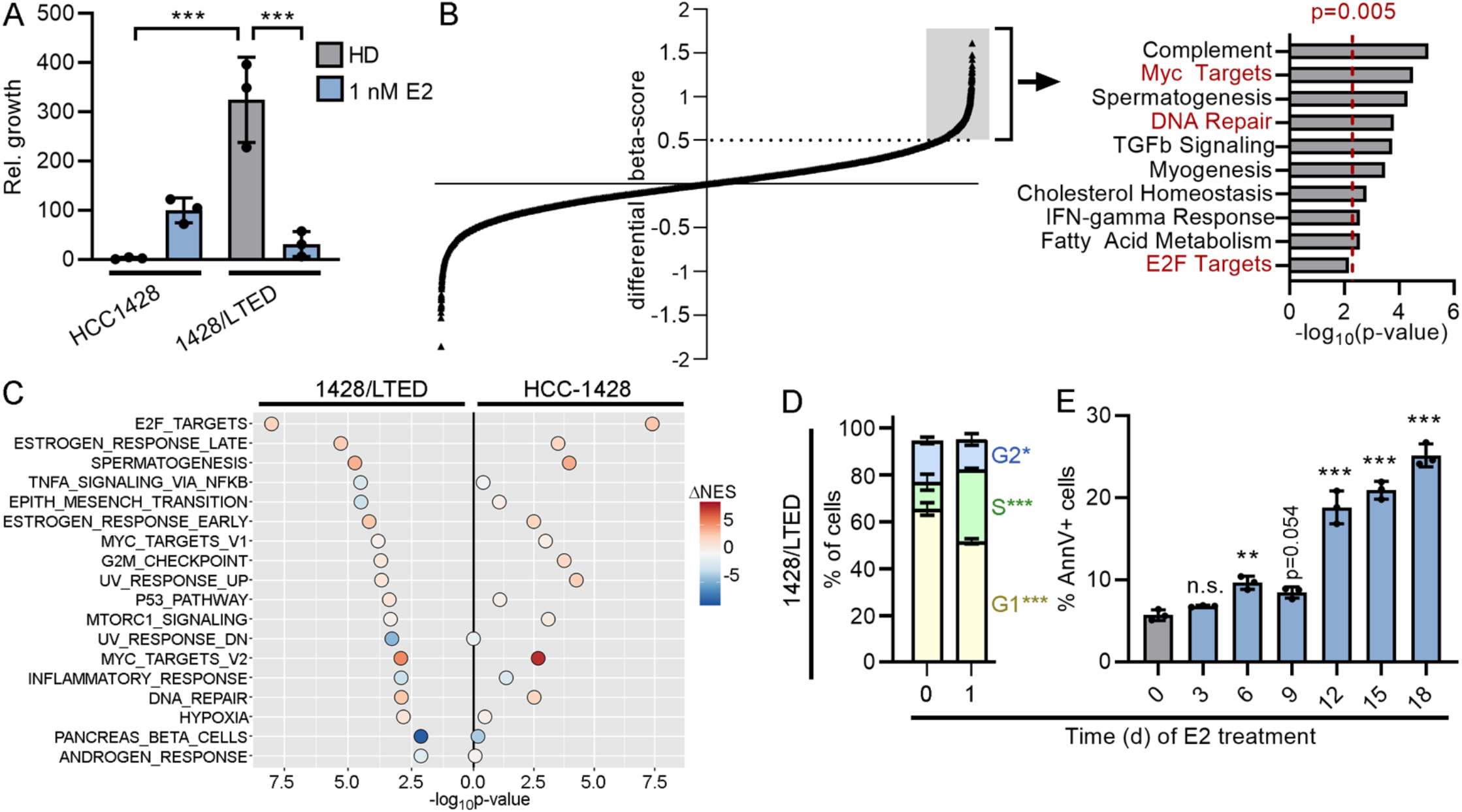
17β-estradiol induces DNA damage in long-term estrogen-deprived cells that therapeutically respond to E2. (A) Parental cells were HD for 3 d prior to seeding. Cells were treated as indicated for 4 wk and relative growth was measured. (B) 1428/LTED cells stably expressing Cas9 were transduced with a sgRNA library. Cells were treated ± 1 nM E2 for 3 wk, and beta-scores were calculated (44). Each point represents one gene. Genes with a differential beta-score ≥0.5 (n=1194, gray box) were analyzed for enrichment with Hallmarks gene sets (*right*). Cell cycle and DNA repair pathways are highlighted in red font. (C) HCC-1428 (HD for 3 d) and 1428/LTED cells were treated ± 1 nM E2 for 24 h, and RNA was harvested for sequencing. Single-sample gene set enrichment analysis (ssGSEA) for Hallmarks pathways was performed for each replicate sample, and normalized gene set enrichment scores (NES) were compared between treatment groups. Gene sets significantly (p<0.05) altered in 1428/LTED cells by E2 treatment are shown, and p-values and differences in mean NES induced by E2 treatment are indicated for each cell line. (D) Cells treated ± 1 nM E2 × 24 h were fixed and stained with propidium iodide (PI). DNA content was measured by flow cytometry. Sub-G1 cells were excluded from plots. Proportions of cells in each phase were compared. (E) Cells were treated ± 1 nM E2 as indicated. Three days after last medium change, cells were harvested, stained with FITC-tagged Annexin V (AnnV), and analyzed by flow cytometry. In (A/D/E), data are shown as mean of triplicates ± SD. *p<0.05, **p<0.005, ***p<0.0005, n.s. = not significant compared to control unless otherwise indicated.

To identify candidate mechanisms underlying therapeutic response to E2, we performed a genome-wide CRISPR/Cas9 knockout screen using 1428/LTED cells. Beta-scores were calculated as a measure of gene essentiality in the presence and absence of E2. We then calculated the difference in beta-scores between E2-treated and HD conditions to identify genes that were required for the growth-inhibitory effect of E2 (i.e., genes that, when lost, rescued from E2; differential beta-score ≥0.5, Fig. 1B). Pathways analysis of this subset of genes revealed enrichment for pathways involved in DNA repair and cell cycle progression (i.e., E2F and Myc targets). Additionally, we observed that knockout of multiple CDKs (*CDK4, CDK9, CDK7, CDK2*) each rescued 1428/LTED cells from the growth inhibitory effects of E2 (Suppl. Table S1). Gene set enrichment analysis of transcriptomic profiles of 1428/LTED cells showed E2-induced engagement of pathways involved in cell cycle (E2F and Myc targets) and DNA damage response (G2/M checkpoint, p53 pathway, UV response, DNA repair), some of which were not detected in parental HCC-1428 cells (Fig. 1C and Suppl. Table S2). In addition, the magnitude of gene expression changes induced by E2 was often greater in 1428/LTED cells than HCC-1428 cells (Suppl. Fig. S1). Consistent with these results, 1428/LTED cells exhibited an accumulation of cells in S-phase following 1 d of E2 treatment, which preceded the onset of apoptosis (Fig. 1D/E). Based on these collective data, we hypothesized that estrogen therapy induces DNA damage during replication.

### 17β-estradiol treatment induces ER- and cell cycle-dependent DNA damage

Cellular response to DNA damage was assessed in HCC-1428 and 1428/LTED cells by immunofluorescent staining for phospho-histone H2AX_Ser139_ (γH2AX), a marker of cellular response to double-strand DNA breaks. 1428/LTED cells exhibited significant increases in γH2AX foci following both 1 and 24 h of E2 treatment, while parental HCC-1428 cells did not (Fig. 2A/B). As another marker of response to DNA breaks, 1428/LTED cells also exhibited an increase in p53 binding protein (53BP1) foci upon treatment with E2 (Suppl. Fig. S2A). BrdU pulse labeling revealed that the majority of cells incurring E2-induced DNA damage (γH2AX+) were in S-phase (BrdU+), and blocking G1-to-S cell cycle progression through treatment with the CDK4/6-selective inhibitor abemaciclib prevented DNA damage induced by E2 (Fig. 2C/D and Suppl. Fig. 3A).

**Figure 2.**
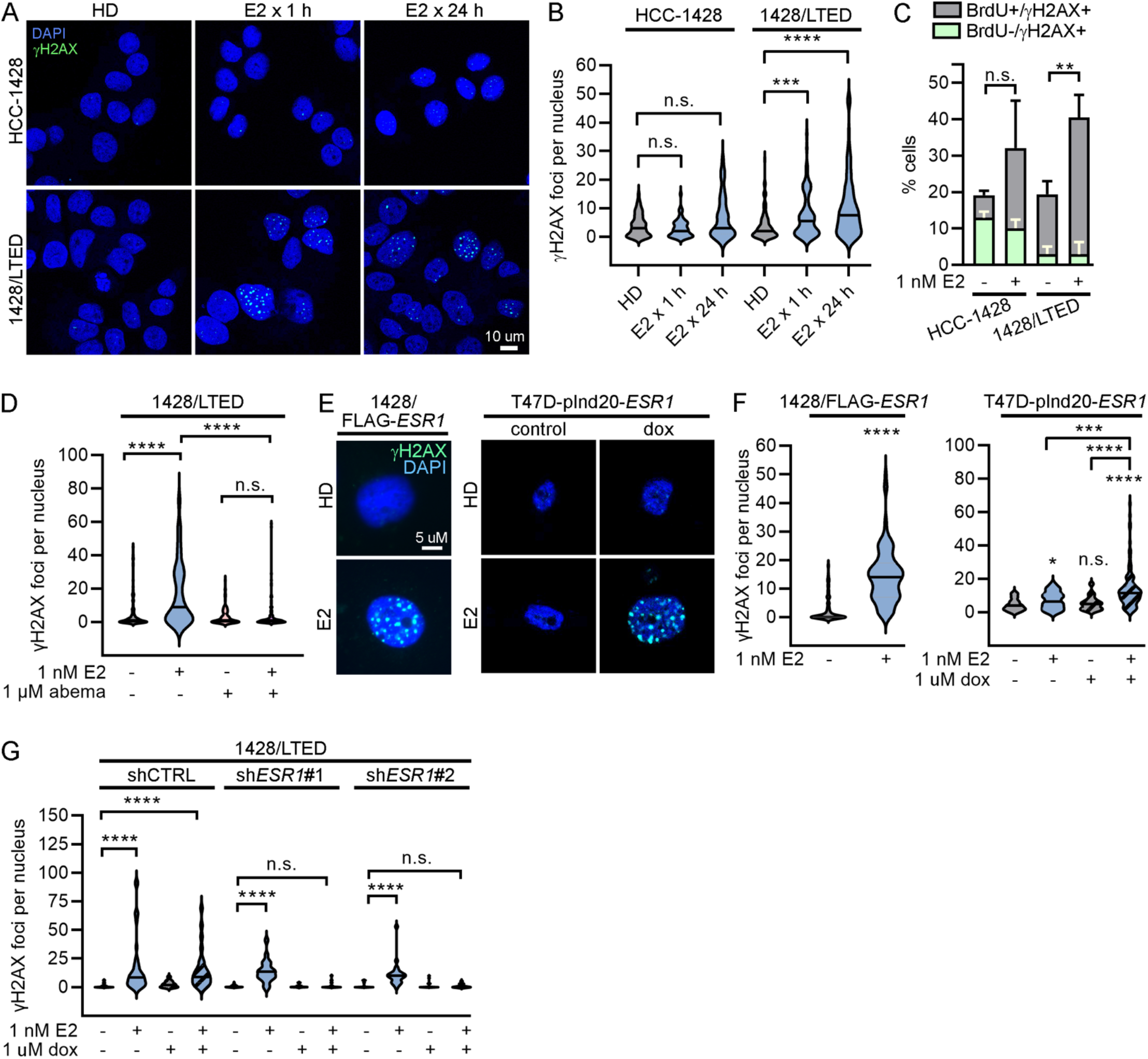
17β-estradiol-induced DNA damage is dependent upon overexpression of ER. (A/B) HCC-1428 (HD x 3 d) and 1428/LTED cells were treated ± 1 nM E2. Cells were fixed and stained for γH2AX (green) and DAPI (blue). γH2AX foci were counted in ≥100 nuclei/group. (C) HCC-1428 (HD × 3 d) and 1428/LTED cells were treated ± E2 for 21 h, and labeled with BrdU for another 3 h. Cells were stained for γH2AX, BrdU, and cleaved PARP for flow cytometry analysis. Cleaved PARP-positive cells were excluded from analysis. Proportions of cells with DNA breaks (γH2AX-positive) that were or were not in S-phase (i.e., did or did not incorporate BrdU) were plotted. Proportions of γH2AX+/BrdU+ cells were statistically compared. Data are shown as mean of triplicates + SD. (D) Cells were treated ± E2 ± abemaciclib × 24 h and analyzed as in (B). (E/F) T47D/pInd20-*ESR1* cells were pretreated with HD × 7 d, and then treated with HD ± dox × 14 d prior to seeding. All cells were then treated as indicated × 24 h and analyzed as in (B). (G) 1428/LTED cell lines expressing dox-inducible shRNA targeting *ESR1* (two independent constructs) or non-silencing control were treated ± dox for 2 d, and then treated ± E2 × 24 h and analyzed as in (B). In (A/E), representative images are shown. *p<0.05, **p<0.005, ***p<0.0005, ****p<0.0001, n.s. = not significant.

1428/LTED cells express increased ER levels compared to parental cells, which we previously demonstrated drives therapeutic response to E2. Forced overexpression of ER (*ESR1*) in HCC-1428 or MDA-MB-415 ER+/HER2-breast cancer cells, and doxycycline (dox)-inducible overexpression of ER in T47D cells drive hormone-independent growth and convert E2 from a growth promoter to a growth suppressor [Suppl. Fig. S4A/B and ref. (12)]. Similar to 1428/LTED cells, exogenous ER overexpression increased E2-induced DNA damage as measured by γH2AX and 53BP1 foci (Fig. 2E/F and Suppl. Fig. S2B/C). Conversely, doxycycline-induced RNAi-mediated knock-down of ER blocked the ability of E2 to induce DNA damage in 1428/LTED cells (Fig. 2G and Suppl. Fig. S4C). ER-negative BT-20 cells did not exhibit an increase in DNA damage following E2 treatment (Suppl. Fig. S5). These data show that ER drives estrogen-inducible DNA damage in ER+ breast cancer cells. Similar to 1428/LTED cells, E2 induced DNA damage in ER-overexpressing cells mainly during S-phase, and treatment with abemaciclib prevents DNA damage upon E2 treatment (Suppl. Fig. S3). Although MCF-7/LTED cells are also growth-inhibited by E2 (15), a DNA damage response was not consistently observed (data not shown), suggesting that this model responds to E2 via a different mechanism. Together, these results indicate that E2 treatment induces DNA damage during replication in ER-overexpressing models that are growth-inhibited by E2.

### Estrogen therapy induces a DNA damage response in human tumors and PDX models

Paired baseline and on-treatment metastatic tumor biopsy specimens were obtained from 2 subjects in clinical trial NCT02188745, which evaluated the therapeutic effects of E2 in patients with endocrine-resistant breast cancer. Both tumors were ER+/PR+ by IHC (Suppl. Fig. S6), and HER2-by FISH (data not shown). Following 2 wk of treatment with E2 (2 mg p.o. TID), γH2AX staining significantly increased (Fig. 3A and Suppl. Fig. S7). ER levels decreased during E2 treatment (Fig. 3A), consistent with ER protein turnover following estrogen-induced ER activation (16). These data suggest that estrogen therapy induces DNA damage in human ER+/HER2-breast tumors.

**Figure 3.**
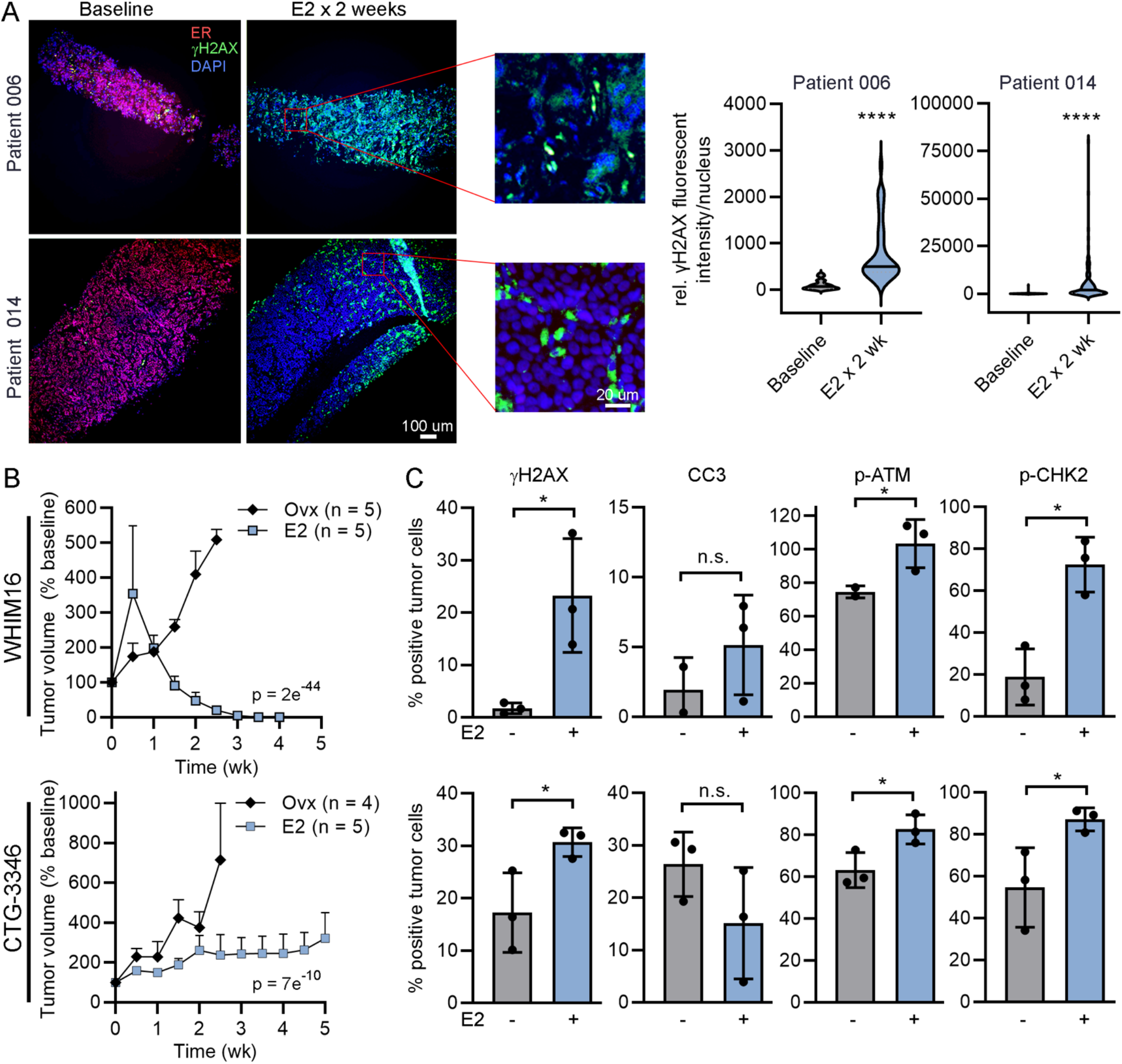
17β-estradiol therapy induces DNA damage in endocrine-resistant tumors. (A) Biopsy samples of advanced ER+ breast tumors were obtained from 2 patients before and 2 wk after treatment with E2. FFPE tumor sections were stained for ER (red), γH2AX (green), and DAPI (blue). γH2AX intensity was quantified in ≥100 nuclei/specimen. Representative exposure-matched image pairs are shown. (B) Ovx mice bearing tumors ∼200 mm^3^ were randomized to treatment ± E2. Tumor volumes were serially measured. Data are shown as mean + SEM, and were analyzed by nonlinear mixed modeling. (C) Tumors (n=3/group) were harvested from ovx mice treated ± E2 for 24 h. FFPE sections were analyzed by IHC. Data are shown as mean ± SD. *p<0.05, ****p<0.0001, n.s. = not significant.

The ability of E2 to induce a DNA damage response was evaluated in 2 estrogen-independent PDX models that are growth-inhibited by E2. The ER+/PR+/HER2-WHIM16 PDX model grows in ovariectomized (ovx) mice, modeling resistance to aromatase inhibitor-induced estrogen deprivation in patients, and regresses upon treatment with E2 [Fig. 3B, Suppl. Fig. S8, and refs. (12,13,15)]. CTG-3346 is a novel PDX model that grows in ovx mice and is growth-inhibited by treatment with E2 (Fig. 3B and Suppl. Fig. S8). CTG-3346 was derived from a patient treated as described as in Suppl. Fig. S9A, and this model retains the ER+/PR+/HER2-status of the patient’s recurrent tumor (Suppl. Fig. S9B and data not shown). CTG-3346 tumors exhibit mutation and copy number loss of *RB1*, and consequent loss of Rb protein as a major tumor suppressor and effector of CDK4/6 (Suppl. Fig. S9C and data not shown), which is consistent with the patient’s tumor resistance to the CDK4/6 inhibitor palbociclib [Suppl. Fig. S9A and ref. (17)]. Despite prior treatment of the patient with multiple lines of endocrine therapies, CTG-3346 retains functional ER, as measured by an increase in mRNA expression of the ER target gene *TFF1* upon treatment of mice with E2 (Suppl. Fig. S9D).

E2 treatment for 24 h significantly increased γH2AX in both PDX models growing in ovx mice (Fig. 3C and Suppl. Fig. S10). However, E2 did not alter cleaved caspase-3 positivity, indicating that the DNA damage reflected by γH2AX is not related to apoptosis. E2 treatment also significantly increased proportions of cells expressing phospho-ATM_S1981_ and phospho-CHK2_T68_, which are markers of an activated DNA damage response (Fig. 3C and Suppl. Fig. S10).

### Estrogen therapy induces transcriptional stress leading to DNA damage

Since E2 induced an increase in S-phase cells, E2-induced DNA damage was dependent upon G1-to-S progression (Figs. 1D and 2D, and Suppl. Fig. S3), and LTED and ER-overexpressing cells exhibit E2-induced hyperactivation of ER transcriptional activity (12,15), we hypothesized that estrogen-induced DNA damage results from ER transcription-driven replication stress via R-loops. While E2 increased replication stress in 1428/LTED and ER-overexpressing cells as shown by proliferating cell nuclear antigen (PCNA) focus formation (18) (Suppl. Fig. S11), cells stimulated to enter S-phase may be expected to undergo replication stress. We there evaluated the contribution of E2-induced transcription to DNA damage.

R-loops are 3-stranded DNA:RNA hybrid structures that form when nascent mRNA re-anneals to the template strand of DNA, impairing re-annealing of complementary DNA strands. While R-loops play roles in transcriptional regulation (19-21), these structures can induce genome instability and DNA breaks, potentially through collision with replication forks (22). Furthermore, ER transcriptional activation has been linked to R-loop formation in MCF-7 cells (23). Using the R-loop structure-specific S9.6 antibody, we observed E2-induced increases in nuclear R-loop formation in 1428/LTED cells that were significantly higher than those detected in parental HCC-1428 cells (Fig. 4A/B). Similarly, 1428/FLAG-*ESR1* and T47D/pInd20-*ESR1* cells showed E2-induced R-loop formation, which was significantly higher upon dox-induced ER overexpression in the T47D model (Suppl. Fig. S12A/B). In addition, IHC analysis of PDX models showed increased R-loops after 24 h of E2 treatment *in vivo* (Fig. 4C). RNase H1 can degrade the RNA specifically within RNA-DNA hybrids. Transient ectopic expression of RNase H1 suppressed E2-induced R-loop formation and prevented DNA damage (Fig.4D/E and Suppl. Fig. S12C), supporting a causative role for R-loops in DNA damage incurred from E2.

**Figure 4.**
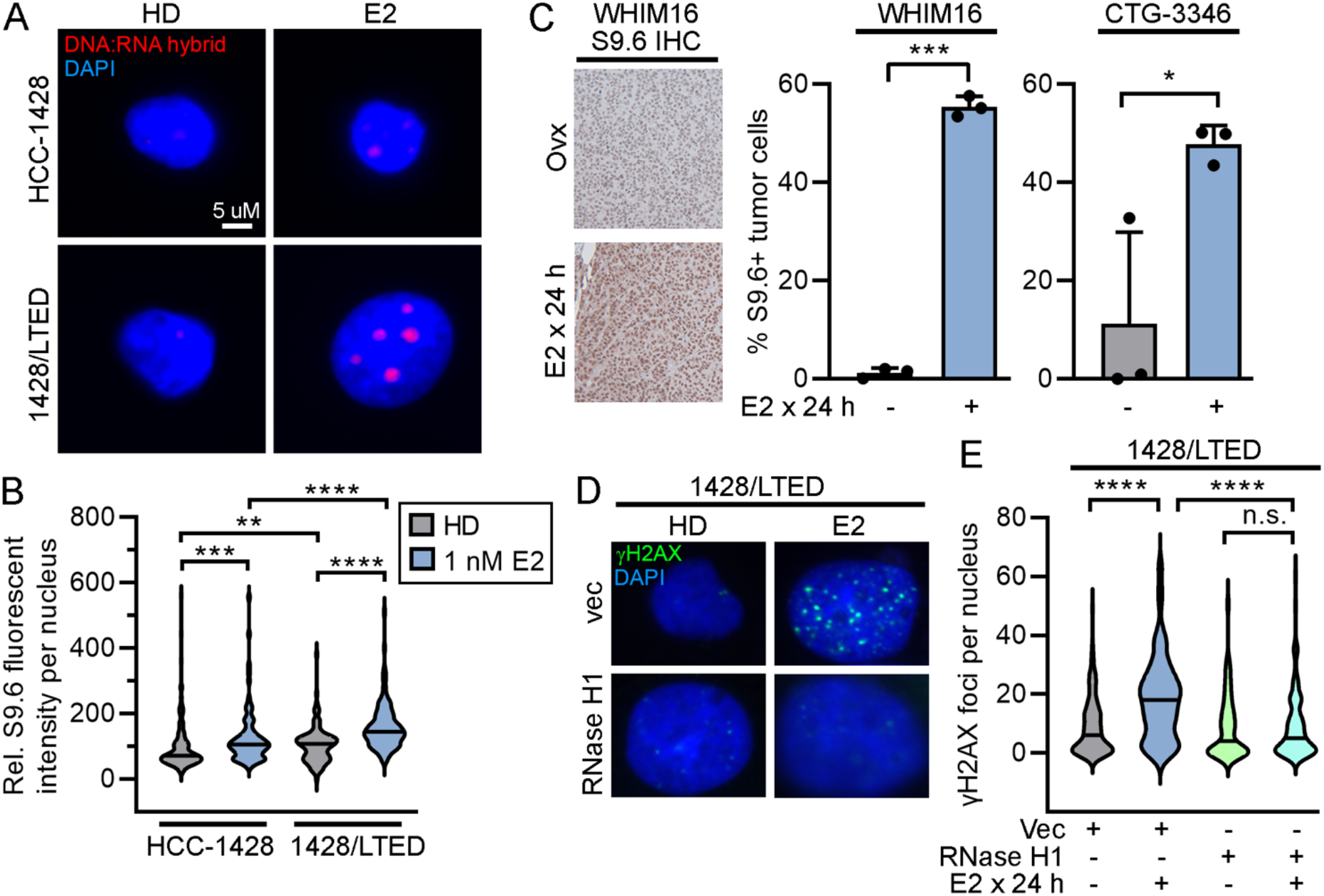
17β-estradiol-induced R-loop formation drives DNA damage. (A/B) Cells were treated ± 1 nM E2 × 24 h, fixed, and stained for DNA/RNA hybrids (S9.6 antibody) and with DAPI. Fluorescence intensity was quantified in ≥100 nuclei/group. (C) Tumors were harvested from ovx mice treated ± E2 (n=3/group). FFPE sections were stained for DNA/RNA hybrids. Proportions of positively staining nuclei were calculated. Data are shown as mean + SD. (D/E) Cells were transiently transfected with plasmids encoding RNase H1 or vector control. Two days later, cells were treated ± 1 nM E2 × 24 h, then fixed and stained for γH2AX and DAPI. γH2AX foci were counted in ≥100 nuclei/group. In (A/D), representative images are shown. *p<0.05, **p<0.005, ***p<0.0005, ****p<0.0001, n.s. = not significant.

### PARP inhibition enhances 17β-estradiol/ER-induced DNA damage and synergistically inhibits growth

We next sought to determine whether inhibiting repair of DNA damage could enhance the therapeutic effects of E2. PARP1/2 are involved in DNA damage repair, and PARP1 play a role in the repair of R-loops and R-loop-associated DNA damage (24,25). We thus hypothesized that inhibition of PARP would enhance E2/ER-induced DNA damage. Treatment with the PARP1/2 inhibitor olaparib synergized with E2 to suppress the growth of 1428/LTED and dox-induced T47/pInd20-*ESR1* cells (Fig. 5A). Accordingly, olaparib potentiated the DNA damage induced by E2 (Fig. 5B/C). These effects were observed in both the *BRCA1*/*2*-wild-type T47D model and the *BRCA2*-altered HCC-1428 models. Olaparib also enhanced R-loop formation when combined with E2 in 1428/LTED and 1428/FLAG-*ESR1* cells (Suppl. Fig. S13).

Currently, PARP inhibitors are FDA-approved for the treatment of breast cancer patients with germline genetic alterations in *BRCA1* or *BRCA2. BRCA1*/*2* encode proteins involved in homologous recombination repair of DNA (26,27) and are postulated to be synthetically lethal with PARP inhibition (28,29). In accordance with the *BRCA1/2* mutation status of these cell lines, HCC-1428 derivatives responded to olaparib treatment as a monotherapy, while T47D derivatives did not (Fig. 5A). However, the synergistic effect of olaparib and E2 in both models indicates that this drug combination is effective regardless of *BRCA1/2* status.

**Figure 5.**
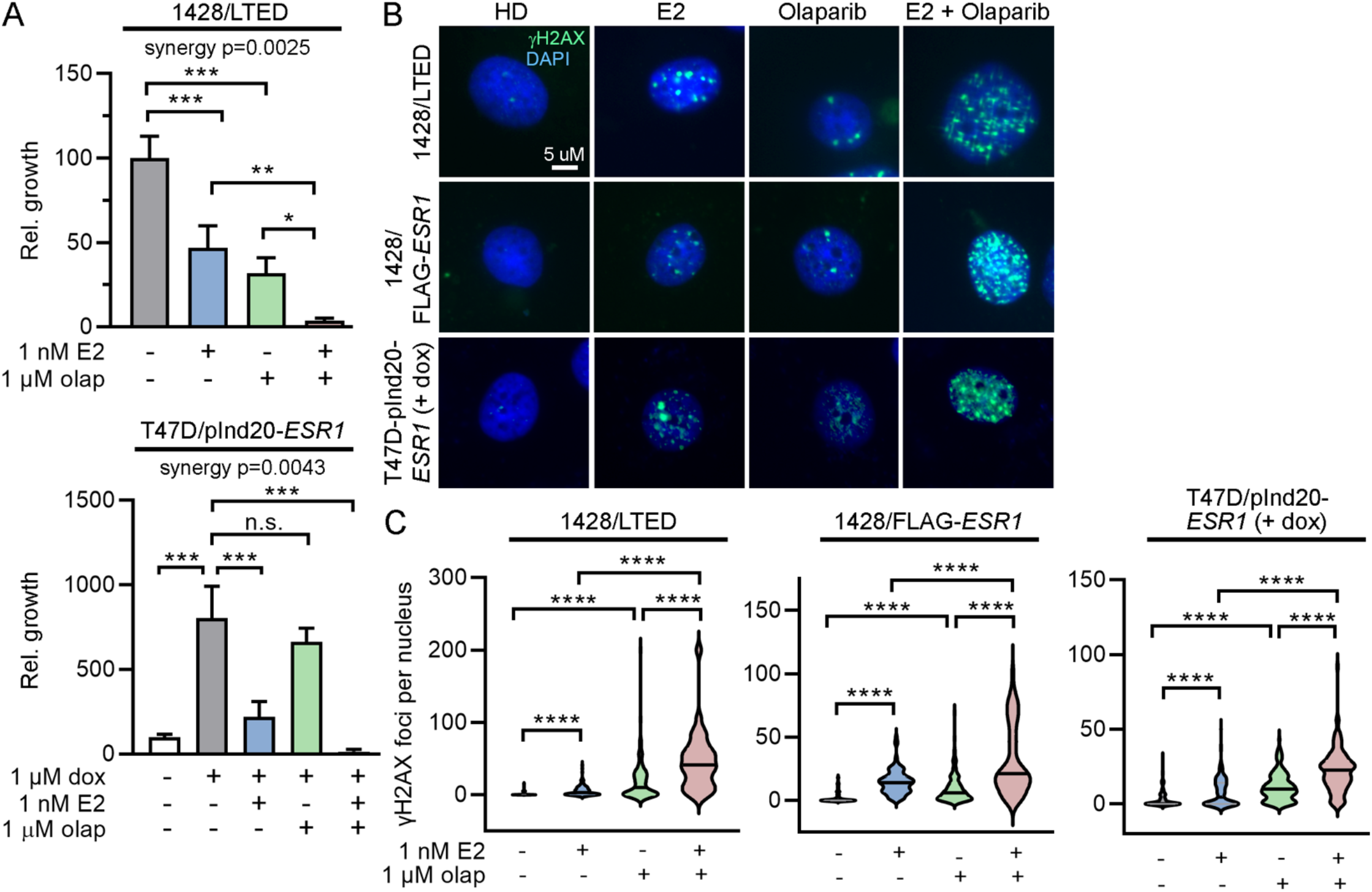
PARP inhibition synergizes with 17β-estradiol to induce DNA damage and inhibit growth *in vitro*. (A) T47D/pInd20-*ESR1* cells were HD × 7 d, and pretreated with HD + dox × 14 d prior to seeding. Both lines were then treated ± olaparib for 2 d, followed by treatment ± olaparib ± E2 for 4 wk, and relative growth was measured. Data are shown as mean of triplicates + SD. (B/C) T47D/pInd20-*ESR1* cells were pretreated as in (A) before seeding. All lines were then treated ± olaparib for 2 d, followed by treatment ± E2 ± olaparib for 24 h. Cells were fixed and stained for γH2AX and DAPI. Representative images are shown. γH2AX foci were counted in ≥100 nuclei/group. *p<0.05, **p<0.005, ***p<0.0005, ****p<0.0001, n.s. = not significant.

### PARP inhibition synergizes with 17β-estradiol to inhibit to tumor growth and prevent recurrence

Ovx mice bearing WHIM16 or CTG-3346 tumors were randomized to treatment with vehicle, olaparib, E2, or the combination of E2 and olaparib. E2 was administered continuously, and olaparib was administered for 28 d (gray shading in Fig. 6A/B). Neither PDX model responded to olaparib monotherapy (Fig. 6A/B and Suppl. Fig. S14). However, olaparib synergized with E2 to delay recurrence of WHIM16 tumors (Fig. 6A/C) and prevent growth of CTG-3346 tumors (Fig. 6B).

**Figure 6.**
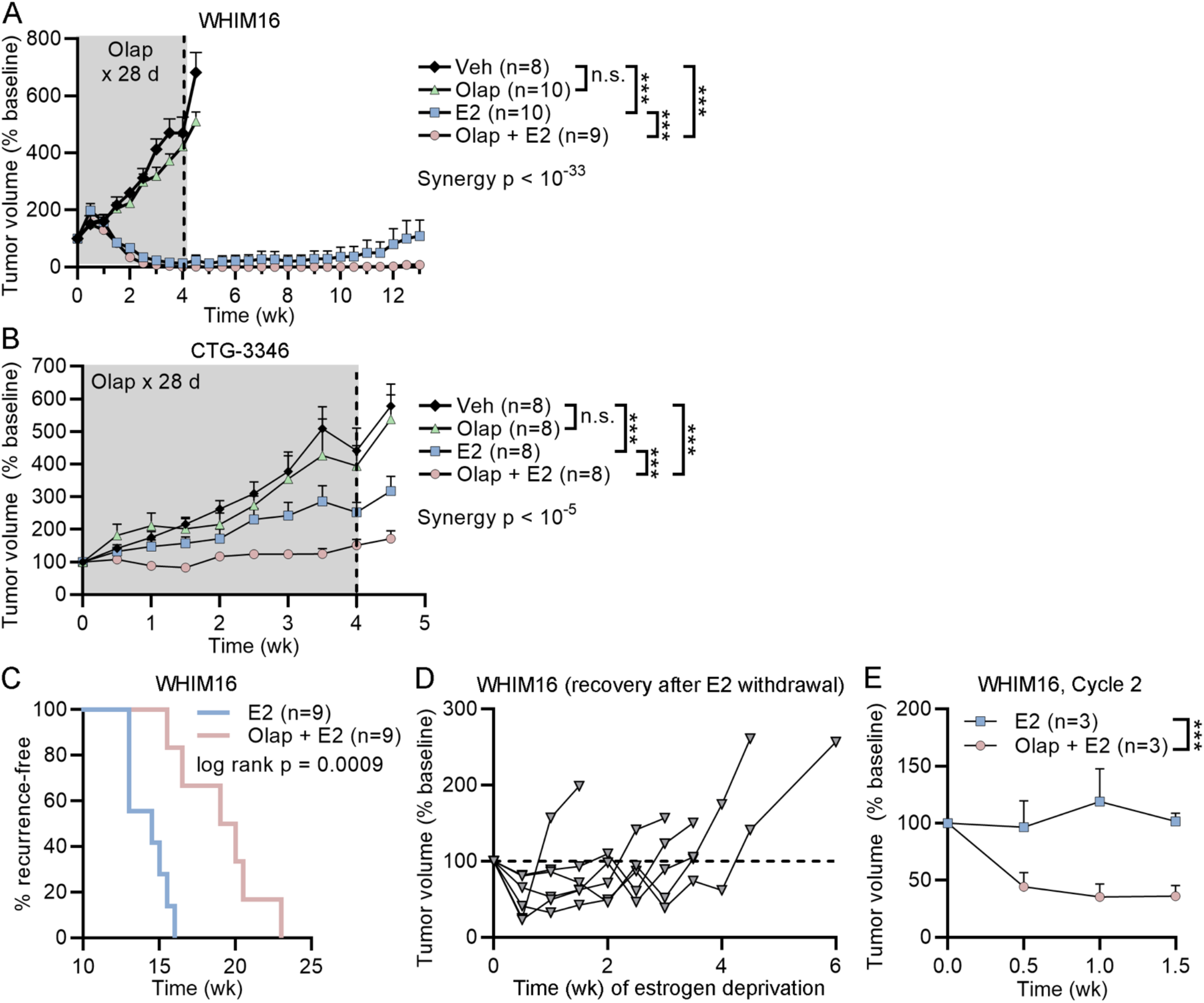
PARP inhibition synergizes with 17β-estradiol against tumors *in vivo* and prevents recurrence. (A/B) Ovx mice bearing tumors ∼200 mm^3^ were randomized to treatment as indicated. E2 was delivered continuously, and olaparib was administered daily for 28 d (gray shading). Data are shown as mean + SEM. (C) After 10 wk of treatment in (A), mice without palpable tumors were monitored for recurrence. Time to recurrence was calculated as time from treatment start until tumors re-grew to baseline volume. Proportions of mice that were recurrence-free over time are shown. (D) Mice bearing tumors that recurred during E2 monotherapy in (C) were treated with estrogen deprivation starting on Day 0, which stunted tumor growth. Each line represents one mouse. Tumors eventually resumed growth, which was defined as two consecutive biweekly volume measurements above baseline. (E) Mice with tumors that resumed estrogen-independent growth in (D) were randomized to treatment with a second cycle of E2 ± olaparib. Data are shown as mean + SEM. ***p<0.0005, n.s. = not significant.

We previously demonstrated that WHIM16 tumors that acquire resistance to E2 treatment are sensitive to estrogen deprivation, and such tumors ultimately regain estrogen-independent growth and re-sensitization to E2 therapy (12). Following recurrence on E2 monotherapy, mice bearing WHIM16 tumors were estrogen-deprived until tumors resumed growth (Fig. 6D). These tumor-bearing mice were then randomized to a second round of treatment with E2 ± olaparib. The addition of olaparib significantly improved the anti-tumor effects of E2 (Fig. 6E and Suppl. Fig. S15). These data collectively suggest that PARP inhibition enhanced the therapeutic effects of E2 by increasing R-loops and DNA damage, and that this synergistic effect can occur in the absence of *BRCA1/2* alterations.

## Discussion

Prior work supported a link between overexpression/amplification of ER, hyperactivation of ER-driven transcription, and therapeutic response to E2 (12,13,15,30). However, the molecular mechanism underlying these effects remained unclear. Herein, we demonstrate that E2 induces ER-dependent S-phase-specific DNA damage and R-loop accumulation that is exacerbated by ER overexpression and adaptation to growth in hormone-depleted conditions. While estrogens have been shown to induce DNA damage and genotoxic stress through several mechanisms, we present evidence of E2-induced DNA damage specifically associated with ER activation and a downstream growth-inhibitory response. This mechanism led us to develop a novel combination therapy with PARP inhibitors and E2, which synergize to enhance therapeutic response in models of endocrine-resistant ER+ breast cancer.

Estrogens can induce DNA damage through both ER-dependent and -independent mechanisms. Estrogens and metabolites have been shown to directly induce genotoxic stress in the absence of ER, including through the formation of DNA adducts (31-33). In our models, E2 induced DNA damage most prominently in LTED and ER-overexpressing models, suggesting an ER-dependent mechanism of DNA damage (Figs. 1B/F/G and 3C/D, and Suppl. Figs. S2 and S5). ER-initiated transcription can engage topoisomerase II? to induce transient double-stranded DNA breaks (34), which are postulated to relieve topological constraints to improve accessibility of DNA for transcription. E2-activated ER can also induce the formation of R-loops leading to DNA breaks at ER-responsive genes (23), but this effect had not previously been associated with a growth-inhibitory response. While previous work suggested a role for R-loops in E2-induced DNA damage, such effects were observed primarily with supraphysiological E2 concentrations of 10-100 nM (23). We utilized 1 nM E2, which is within the pre-menopausal serum physiological range in humans and achievable pharmacologically in serum through oral E2 treatment in post-menopausal patients (7,35).

Using the clinically relevant dose of 1 nM E2, we observed accumulation of R-loops that were significantly more abundant in ER-overexpressing cells that are growth-inhibited by E2 (Fig. 4B and Suppl. Fig. S12A/B).

Similar to estrogen therapy for endocrine-resistant breast cancer, a subset of patients with castration-resistant prostate cancer benefit from treatment with androgens, and this effect may be dependent upon androgen receptor overexpression or amplification. Androgen treatment induces DNA damage, which is exacerbated in *BRCA2*-deficient cells (36,37). A recent clinical study showed that 16/36 (44%) patients with metastatic castration-resistant prostate cancer experienced a PSA_50_ response (i.e., a decline ≥50% from baseline in plasma prostate-specific antigen levels) upon treatment with bipolar androgen therapy (BAT; testosterone cypionate/enanthate administered every 28 d) in combination with olaparib regardless of tumor *BRCA1/2* status (38). Our results do not indicate that *BRCA1/2* mutations are a requirement for therapeutic response to E2; however, *BRCA1/2* mutations or other defects in homologous recombination-mediated DNA repair may increase the likelihood that tumors therapeutically respond to estrogen by preventing repair of estrogen/ER-induced DNA damage. In this study, we leveraged combination treatment with a PARP inhibitor due to the roles of PARP1/2 in repair of R-loops and DNA damage. However, other inhibitors of the DNA damage response may also synergize with estrogen therapy; as new inhibitors enter clinic development (e.g., agents targeting WEE1, ATR, etc.), this concept will warrant further investigation.

This work raises several points surrounding clinical implementation of estrogen therapy. Firstly, these findings offer the possibility of expanding the use of PARP inhibitors to a larger patient population. Currently, PARP inhibitors are FDA-approved for breast cancer only in the setting of germline *BRCA1/2* genetic alterations, which constitute ∼5% of breast cancer cases (39). In our studies, olaparib synergized with E2 in both *BRCA1/2*-mutant and -wild-type models, and synergy was seen even in models that were non-responsive to olaparib monotherapy (Figs. 5A and 6A/B). A clinical indication for E2/PARP inhibitor combination therapy in advanced ER+/HER2-disease regardless of *BRCA1/2* status would substantially increase the number of patients who could benefit from a repurposed PARP inhibitor.

Another point raised by this work is the efficacy of E2 therapy in the landscape of CDK4/6 inhibitors. Treatment with CDK4/6 inhibitors has become a standard of care in advanced ER+ breast cancer, and acquired resistance to CDK4/6 inhibitors is an emerging clinical problem. Since CDK4/6 inhibitors are a relatively recent development in the field, strategies for the treatment of ER+ breast cancer following progression on CDK4/6 inhibitors remain underexplored. We developed the CTG-3346 PDX tumor from a patient with acquired resistance to the CDK4/6 inhibitor palbociclib. CTG-3346 exhibits loss/mutation of Rb (Suppl. Fig. S9A/C), consistent with acquired resistance to CDK4/6 inhibition (17). The growth-inhibitory effects of E2 in this model (Fig. 3B) indicate that E2 therapy remains an effective treatment option in the post-CDK4/6 inhibitor setting.

Although E2 therapy may remain effective despite resistance to CDK4/6 inhibitors, this work cautions against combining E2 with CDK4/6 inhibitors. Our data indicate that cell cycle progression and DNA replication are required for E2-induced DNA damage, and CDK4/6 inhibition antagonized the DNA-damaging effects of E2 (Fig. 2D and Suppl. Fig. S3C). A similar antagonistic effect has been reported with CDK4/6 inhibitors co-administered with cytotoxic chemotherapies that target S-phase and mitotic cells (40,41). However, the antagonistic effects of these drug combinations may be dependent upon the order and timing with which they are administered. Several studies have suggested that pre-treatment with cytotoxic chemotherapies followed by treatment with CDK4/6 inhibitors elicits synergistic effects. Such synergy may be due to the repressed transcription of DNA repair genes by CDK4/6 inhibitors, which could impede DNA repair following treatment with cytotoxic chemotherapy (42,43). If E2 is considered a DNA-damaging agent in endocrine-resistant ER+ breast cancer, then strategically timed CDK4/6 inhibition following E2 treatment may convert these seemingly antagonistic drugs to synergistic; this concept warrants further study.

## Acknowledgements

This work was supported by Susan G. Komen (CCR1533084 to TWM) and NIH (R01CA200994, R01CA267691, R01CA262232, R01CA211869 to TWM, F31CA243409 to NAT, the Center for Quantitative Biology at Dartmouth P20GM130454, Dartmouth College Cancer Center Support Grant P30CA023108). We thank Alan Eastman, Chris Shoemaker, Scott Kaufmann, and Matthew Goetz for helpful discussions, and the following Dartmouth Cancer Center Shared Resources for their support: Mouse Modeling; Pathology; Biostatistics; Bioinformatics; Microscopy; Genomics & Molecular Biology.

## References Cited

1. Shiino S, Kinoshita T, Yoshida M, Jimbo K, Asaga S, Takayama S, et al. Prognostic Impact of Discordance in Hormone Receptor Status Between Primary and Recurrent Sites in Patients With Recurrent Breast Cancer. Clinical breast cancer 2016;16(4):e133–40.

2. Osborne CK, Pippen J, Jones SE, Parker LM, Ellis M, Come S, et al. Double-blind, randomized trial comparing the efficacy and tolerability of fulvestrant versus anastrozole in postmenopausal women with advanced breast cancer progressing on prior endocrine therapy: results of a North American trial. J Clin Oncol 2002;20(16):3386–95.

3. Haddow A, Watkinson JM, Paterson E, Koller PC. Influence of Synthetic Oestrogens on Advanced Malignant Disease. British medical journal 1944;2(4368):393–8.

4. Ingle JN, Ahmann DL, Green SJ, Edmonson JH, Bisel HF, Kvols LK, et al. Randomized clinical trial of diethylstilbestrol versus tamoxifen in postmenopausal women with advanced breast cancer. The New England journal of medicine 1981;304(1):16–21.

5. Peethambaram PP, Ingle JN, Suman VJ, Hartmann LC, Loprinzi CL. Randomized trial of diethylstilbestrol vs. tamoxifen in postmenopausal women with metastatic breast cancer. An updated analysis. Breast Cancer Res Treat 1999;54(2):117–22.

6. Kennedy BJ. Systemic Effects of Androgenic and Estrogenic Hormones in Advanced Breast Cancer. J Am Geriatr Soc 1965;13:230–5.

7. Ellis MJ, Gao F, Dehdashti F, Jeffe DB, Marcom PK, Carey LA, et al. Lower-dose vs high-dose oral estradiol therapy of hormone receptor-positive, aromatase inhibitor-resistant advanced breast cancer: a phase 2 randomized study. Jama 2009;302(7):774–80.

8. Iwase H, Yamamoto Y, Yamamoto-Ibusuki M, Murakami KI, Okumura Y, Tomita S, et al. Ethinylestradiol is beneficial for postmenopausal patients with heavily pre-treated metastatic breast cancer after prior aromatase inhibitor treatment: a prospective study. British journal of cancer 2013;109(6):1537–42.

9. Zucchini G, Armstrong AC, Wardley AM, Wilson G, Misra V, Seif M, et al. A phase II trial of low-dose estradiol in postmenopausal women with advanced breast cancer and acquired resistance to aromatase inhibition. European journal of cancer 2015;51(18):2725–31.

10. Lonning PE, Taylor PD, Anker G, Iddon J, Wie L, Jorgensen LM, et al. High-dose estrogen treatment in postmenopausal breast cancer patients heavily exposed to endocrine therapy. Breast Cancer Res Treat 2001;67(2):111–6.

11. Agrawal A, Robertson JF, Cheung KL. Efficacy and tolerability of high dose “ethinylestradiol” in post-menopausal advanced breast cancer patients heavily pre-treated with endocrine agents. World journal of surgical oncology 2006;4:44.

12. Traphagen NA, Hosford SR, Jiang A, Marotti JD, Brauer BL, Demidenko E, et al. High estrogen receptor alpha activation confers resistance to estrogen deprivation and is required for therapeutic response to estrogen in breast cancer. Oncogene 2021;40(19):3408–21.

13. Li S, Shen D, Shao J, Crowder R, Liu W, Prat A, et al. Endocrine-therapy-resistant ESR1 variants revealed by genomic characterization of breast-cancer-derived xenografts. Cell reports 2013;4(6):1116–30.

14. Miller TW, Hennessy BT, Gonzalez-Angulo AM, Fox EM, Mills GB, Chen H, et al. Hyperactivation of phosphatidylinositol-3 kinase promotes escape from hormone dependence in estrogen receptor-positive human breast cancer. J Clin Invest 2010;120(7):2406–13.

15. Hosford SR, Shee K, Wells JD, Traphagen NA, Fields JL, Hampsch RA, et al. Estrogen therapy induces an unfolded protein response to drive cell death in ER+ breast cancer. Molecular oncology 2019;13(8):1778–94.

16. Pakdel F, Le Goff P, Katzenellenbogen BS. An assessment of the role of domain F and PEST sequences in estrogen receptor half-life and bioactivity. The Journal of steroid biochemistry and molecular biology 1993;46(6):663–72.

17. Kumarasamy V, Vail P, Nambiar R, Witkiewicz AK, Knudsen ES. Functional Determinants of Cell Cycle Plasticity and Sensitivity to CDK4/6 Inhibition. Cancer Res 2021;81(5):1347–60.

18. Mailand N, Gibbs-Seymour I, Bekker-Jensen S. Regulation of PCNA-protein interactions for genome stability. Nature reviews Molecular cell biology 2013;14(5):269–82.

19. Ginno PA, Lott PL, Christensen HC, Korf I, Chedin F. R-loop formation is a distinctive characteristic of unmethylated human CpG island promoters. Mol Cell 2012;45(6):814–25.

20. Skourti-Stathaki K, Proudfoot NJ, Gromak N. Human senataxin resolves RNA/DNA hybrids formed at transcriptional pause sites to promote Xrn2-dependent termination. Mol Cell 2011;42(6):794–805.

21. Yang Y, McBride KM, Hensley S, Lu Y, Chedin F, Bedford MT. Arginine methylation facilitates the recruitment of TOP3B to chromatin to prevent R loop accumulation. Mol Cell 2014;53(3):484–97.

22. Hamperl S, Bocek MJ, Saldivar JC, Swigut T, Cimprich KA. Transcription-Replication Conflict Orientation Modulates R-Loop Levels and Activates Distinct DNA Damage Responses. Cell 2017;170(4):774–86 e19.

23. Stork CT, Bocek M, Crossley MP, Sollier J, Sanz LA, Chedin F, et al. Co-transcriptional R-loops are the main cause of estrogen-induced DNA damage. eLife 2016;5.

24. Ye BJ, Kang HJ, Lee-Kwon W, Kwon HM, Choi SY. PARP1-mediated PARylation of TonEBP prevents R-loop-associated DNA damage. DNA Repair (Amst) 2021;104:103132.

25. Cristini A, Groh M, Kristiansen MS, Gromak N. RNA/DNA Hybrid Interactome Identifies DXH9 as a Molecular Player in Transcriptional Termination and R-Loop-Associated DNA Damage. Cell reports 2018;23(6):1891–905.

26. Moynahan ME, Chiu JW, Koller BH, Jasin M. Brca1 controls homology-directed DNA repair. Mol Cell1999;4(4):511–8.

27. Moynahan ME, Pierce AJ, Jasin M. BRCA2 is required for homology-directed repair of chromosomal breaks. Mol Cell 2001;7(2):263–72.

28. Farmer H, McCabe N, Lord CJ, Tutt AN, Johnson DA, Richardson TB, et al. Targeting the DNA repair defect in BRCA mutant cells as a therapeutic strategy. Nature 2005;434(7035):917–21.

29. Bryant HE, Schultz N, Thomas HD, Parker KM, Flower D, Lopez E, et al. Specific killing of BRCA2-deficient tumours with inhibitors of poly(ADP-ribose) polymerase. Nature 2005;434(7035):913–7.

30. Kota K, Brufsky A, Oesterreich S, Lee A. Estradiol as a Targeted, Late-Line Therapy in Metastatic Breast Cancer with Estrogen Receptor Amplification. Cureus 2017;9(7):e1434.

31. Savage KI, Matchett KB, Barros EM, Cooper KM, Irwin GW, Gorski JJ, et al. BRCA1 deficiency exacerbates estrogen-induced DNA damage and genomic instability. Cancer Res 2014;74(10):2773–84.

32. Yue W, Wang JP, Li Y, Fan P, Liu G, Zhang N, et al. Effects of estrogen on breast cancer development: Role of estrogen receptor independent mechanisms. Int J Cancer 2010;127(8):1748–57.

33. Tripathi K, Mani C, Somasagara RR, Clark DW, Ananthapur V, Vinaya K, et al. Detection and evaluation of estrogen DNA-adducts and their carcinogenic effects in cultured human cells using biotinylated estradiol. Mol Carcinog 2017;56(3):1010–20.

34. Ju BG, Lunyak VV, Perissi V, Garcia-Bassets I, Rose DW, Glass CK, et al. A topoisomerase IIbeta-mediated dsDNA break required for regulated transcription. Science 2006;312(5781):1798–802.

35. Kaaks R, Tikk K, Sookthai D, Schock H, Johnson T, Tjonneland A, et al. Premenopausal serum sex hormone levels in relation to breast cancer risk, overall and by hormone receptor status - results from the EPIC cohort. Int J Cancer 2014;134(8):1947–57.

36. Chatterjee P, Schweizer MT, Lucas JM, Coleman I, Nyquist MD, Frank SB, et al. Supraphysiological androgens suppress prostate cancer growth through androgen receptor-mediated DNA damage. J Clin Invest 2019;130:4245–60.

37. Schweizer MT, Antonarakis ES, Wang H, Ajiboye AS, Spitz A, Cao H, et al. Effect of bipolar androgen therapy for asymptomatic men with castration-resistant prostate cancer: results from a pilot clinical study. Sci Transl Med 2015;7(269):269ra2.

38. Schweizer MT, Gulati R, Yezefski T, Cheng HH, Mostaghel E, Haffner MC, et al. Bipolar androgen therapy plus olaparib in men with metastatic castration-resistant prostate cancer. Prostate Cancer Prostatic Dis 2022.

39. Rebbeck TR, Mitra N, Wan F, Sinilnikova OM, Healey S, McGuffog L, et al. Association of type and location of BRCA1 and BRCA2 mutations with risk of breast and ovarian cancer. JAMA 2015;313(13):1347–61.

40. Dean JL, McClendon AK, Knudsen ES. Modification of the DNA damage response by therapeutic CDK4/6 inhibition. J Biol Chem 2012;287(34):29075–87.

41. Pikman Y, Alexe G, Roti G, Conway AS, Furman A, Lee ES, et al. Synergistic Drug Combinations with a CDK4/6 Inhibitor in T-cell Acute Lymphoblastic Leukemia. Clin Cancer Res 2017;23(4):1012–24.

42. Cao J, Zhu Z, Wang H, Nichols TC, Lui GYL, Deng S, et al. Combining CDK4/6 inhibition with taxanes enhances anti-tumor efficacy by sustained impairment of pRB-E2F pathways in squamous cell lung cancer. Oncogene 2019;38(21):4125–41.

43. Salvador-Barbero B, Alvarez-Fernandez M, Zapatero-Solana E, El Bakkali A, Menendez MDC, Lopez-Casas PP, et al. CDK4/6 Inhibitors Impair Recovery from Cytotoxic Chemotherapy in Pancreatic Adenocarcinoma. Cancer Cell 2020;38(4):584.

44. Li W, Koster J, Xu H, Chen CH, Xiao T, Liu JS, et al. Quality control, modeling, and visualization of CRISPR screens with MAGeCK-VISPR. Genome biology 2015;16:281.

