## Supplemental Material for "Estrogen therapy induces receptor-dependent DNA damage enhanced by PARP inhibition in ER+ breast cancer"

**Supplemental Materials for Traphagen et al.**

**Supplemental Methods**

**Cell culture**
HCC-1428 cells were obtained from American Type Culture Collection (ATCC) and cultured in DMEM with 10% FBS (HyClone Laboratories). T47D/pInd20-*ESR1* cells (1) were maintained in DMEM + 10% FBS. HCC-1428/LTED cells were generated through long-term (~1 yr) culture in hormone-depleted conditions (HD) via phenol red-free DMEM with 10% dextran/charcoal-treated FBS (DCC-FBS; Hyclone Laboratories) as described (2). HCC-1428/FLAG-*ESR1* cells were generated through stable integration of FLAG-tagged *ESR1* and maintained in HD medium (ThermoFisher Scientific) as described (1). Cell lines were confirmed to be mycoplasma-free (Universal Mycoplasma Detection Kit; ATCC) and authenticated by STR genotyping at the University of Vermont Cancer Center DNA Analysis Facility.

**Immunohistochemistry**
Five-micron sections of FFPE tissue were mounted on slides, deparaffinized in xylene, and rehydrated in a graded ethanol series. For IHC, antigen retrieval was performed in citrate buffer, pH 6 (VWR), and sections were permeabilized with 0.2% Triton X-100 in PBS. Sections were blocked in 5% goat serum in PBS for 30 min, then incubated overnight in blocking solution with primary antibody at 4°C. Primary antibodies included: phospho-histone $H2AX_{S139}$ (Cell Signaling Technology #9718); S9.6 (Millipore Sigma #MABE1095), cleaved caspase 3 (Cell Signaling Technology #9664); phospho-$ATM_{S1981}$ (Abcam #81292); phospho-$Chk2_{T68}$ (Cell Signaling Technology #2917); progesterone receptor A/B (Cell Signaling Technology #3153). Sections were washed and treated with 0.3% hydrogen peroxide, and signal was developed using the VectaStain Elite ABC-HRP kit with DAB substrate (Vector Laboratories). Sections were counterstained with hematoxylin. IHC staining of PDX tumors was quantified in 3 representative microscopic fields/tumor at 200x magnification using Halo software (Indica Labs).

**Immunofluorescence**
*Tumors*: Slide-mounted 5-micron sections of FFPE tissue were processed as above through the permeabilization step, and blocked in 1% BSA in PBS for 1 h. Sections were incubated with primary antibody in blocking solution overnight at 4°C, washed 3 times, incubated with a fluorescently labeled secondary antibody for 1 h at room temp., washed 3 times, and mounted in ProLong Gold with DAPI (ThermoFisher Scientific). Sections were imaged at 100x magnification, and fluorescent intensity was quantified using CellProfiler software (3).

*Cell lines*: Cells were grown on glass coverslips, treated as indicated in figure legends, and fixed in 1.85% formaldehyde. Cells were permeabilized with 0.1% Triton X-100 in PBS, and blocked in 1% BSA in PBS for 1 h. Cells were incubated with primary antibody in blocking solution overnight at 4°C, wash 3 times, and incubated with a fluorescently labeled secondary antibody at 1 h at room temp. Coverslips were mounted in ProLong Gold with DAPI (ThermoFisher Scientific). Cells were imaged at 400x magnification. Staining was quantified by manually counting γH2AX, 53BP1, and PCNA foci per nucleus, and fluorescent intensity was quantified in S9.6-immunostained cells using CellProfiler software (3). Staining was quantified in ≥100 cells per treatment group.

Primary antibodies included: 53BP1 (Cell Signaling Technology #4937); phospho-histone $H2AX_{S139}$ (Cell Signaling Technology #9718); S9.6 (Millipore Sigma #MABE1095); estrogen receptor alpha (Santa Cruz Biotechnology #sc-8002); PCNA (Cell Signaling Technology #8580). Secondary antibodies included: AlexaFluor 488 goat anti-rabbit IgG (Life Technologies #A11034); AlexaFluor 647 goat anti-mouse IgG Fab2 (Cell Signaling Technology #4410).

**Immunoblotting**
All chemicals were purchased from Sigma unless otherwise noted. Cells were lysed in RIPA buffer (20 mM Tris, pH 7.4, 150 mM NaCl, 1% NP-40, 10% glycerol, 1 mM EDTA, 1 mM EGTA, 5 mM NaPPi, 50 mM NaF, 10 mM Na β-glycerophosphate) plus fresh HALT protease inhibitor cocktail (Pierce) and 1 mM $Na_3VO_4$ (New

England Biolabs). Cell lysates were sonicated at 30% power for 15 s, centrifuged at 17,000 x *g* for 10 min at 4°C, and protein content in supernatant was quantified by BCA Assay (Pierce). Protein extracts were reduced and denatured in LDS sample buffer (GenScript) plus 1.25% β-mercaptoethanol. Proteins were separated by SDS-PAGE, transferred to nitrocellulose, and stained with Ponceau S to visually confirm protein loading and transfer. Blots were probed with primary antibodies against ER (Santa Cruz Biotechnology; cat.# sc-8002), vinculin (Cell Signaling Technology; cat.# 13901), Rb (Cell Signaling Technology; cat.#9309), phospho-$Rb_{S780}$ (Cell Signaling Technology; cat.#9307), and FLAG (Millipore Sigma; cat.# F3165). Signal was detected with DyLight conjugated secondary antibodies (Cell Signaling Technology) using the LI-COR Odyssey system (LI-COR Biosciences).

**Mouse studies and tumor volume statistical analyses**

Studies were approved by the Dartmouth College IACUC. The WHIM16 tumor model was obtained from the Washington University HAMLET core. The CTG-3346 tumor model was developed from recurrent breast tumor tissue obtained by ultrasound-guided biopsy at Dartmouth-Hitchcock Medical Center: fresh tissue cores were placed into transport medium, stored at 4°C, and sent overnight to Champion Oncology. A tissue core was implanted s.c. into two NSG female mice (without supplementation with exogenous E2). Tumor growth was monitored by palpation. After ~8 months, P1 tumors were harvested and cut into fragments that were serially transplanted into P2 ovariectomized (ovx) mice. P4 and P5 tumors were used for treatment studies herein. The CTG-3346 PDX model is available upon request.

Ovx NSG mice were obtained from the Dartmouth Cancer Center Mouse Modeling Shared Resource. Tumor fragments (~8 $mm^3$) were implanted subcutaneously (s.c.) into ovx mice aged 4-6 wk. Tumor dimension were measured twice weekly with calipers, and tumor volume was calculated as: length x $width^2/2$. Mice were randomized to treatment groups when tumor volume reached ~200 $mm^3$. E2 was administered by s.c. beeswax pellet containing 1 mg E2 as described in ref. (4), which were replaced every 30 d. Olaparib (MedChemExpress) was dissolved in 10% (2-hydroxypropyl)-β-cyclodextrin (Sigma-Aldrich) in PBS. Olaparib (50 mg/kg/d) or vehicle was delivered by daily intraperitoneal (i.p.) injection.

Tumor growth data were analyzed using the following linear mixed model: $Log_{10}(\text{tumor volume}_{it}) = a_i + b * t + e_{it}$, where *i* represents the $i^{th}$ mouse, *t* represents time of tumor volume measurement, $a_i$ represents the mouse-specific log tumor volume at $t = 0$, b represents the rate of tumor volume growth, and $e_{it}$ represents deviation of measurements from the model over time (5,6). Mouse heterogeneity (the baseline tumor volume) is represented by variance of $a_i$, and $b * \log_e[10] * 100$ indicates tumor volume increase (%) per week. Treatment groups were compared using a *z*-test for slopes with standard error derived from the output of the function *lme* from the library *nlme* in R. Synergy was determined as described in ref. (7). Comparison of tumor recurrence curves was performed by log-rank test.

**CRISPR/Cas9 knockout screen**

HCC-1428/LTED cells stably expressing Cas9-Blast were generated by lentiviral transduction of the lentiCas9-Blast plasmid (gift from Feng Zhang, Broad Institute; obtained from Addgene #52962). Two days following viral transduction, cells were selected with 5 µg/mL blasticidin for 3 wk, and Cas9 expression was confirmed by RT-qPCR. 120 million cells stably expressing Cas9 were lentivirally transduced with the human Brunello CRISPR knockout pooled library (gift from David Root and John Doench, Broad Institute; obtained from Addgene #73178) at an MOI of 0.3. Two days following viral transduction, cells were treated with 1 µg/mL puromycin for 7 d. Fourteen days following transduction, cells were trypsinized and split into 3 treatment groups at 35 million cells per group. Group A was immediately flash-frozen to represent baseline. Group B was treated with HD medium for 3 wk. Group C was treated with HD medium + 1 nM E2 for 3 wk. Cells in Groups B and C were then trypsinized and flash-frozen. Genomic DNA was collected from the 3 groups using the Blood DNA Maxi Kit (Qiagen). Samples were prepared by PCR amplification of the genomically integrated sgRNA sequences: 100 µg of genomic DNA per sample was split into 100 PCR reactions, using 30 PCR cycles, followed by pooling of PCR products. PCR products were purified with the QIAquick PCR Purification Kit (Qiagen). Amplicon size was confirmed by gel electrophoresis. A secondary PCR was performed with 100 ng per sample for 10 cycles to attach Illumina adapters. PCR products were purified by adding 0.8X and subsequently 2X KAPA Pure beads. Libraries were sequenced using an Illumina NextSeq 500 (2 x 75 bp) on High Output setting. The 5' ends of reads were trimmed to 5'-CACCG-3' using Cutadapt (8). MAGeCK (v.0.5.9) was used to extract read counts for each sgRNA using the count function (9). The mle function was used to compare read counts from cells treated ± E2 after controlling for baseline counts and library size. The output included beta scores. A differential beta score was calculated by taking the difference in beta scores between the E2-treated and HD conditions. To determine significant pathway enrichment, gene set enrichment analysis for Hallmarks pathways was performed using the Molecular Signatures Database (10).

**RNA-seq analysis**

HCC-1428 and HCC-1428/LTED cells were treated with HD medium for 3 d, and then treated ± 1 nM E2 × 24 h in triplicate in 100-mm dishes. RNA was extracted using RNeasy Plus Mini Kit (Qiagen catalog # 1062832). RNA quality was assessed on a fragment analyzer (Advanced Analytical Technologies, Agilent), and RNA was quantified by Qubit. In preparation for RNA sequencing (RNA-seq), ribo-depleted libraries were prepared from 2.5 µg of total RNA using the Globin-Zero Gold (Illumina catalog # GZG1206) and TruSeq Stranded Total RNA (Illumina catalog # RS-122-2201) workflows according to manufacturer's instructions. Each library was uniquely barcoded, quantified by qPCR (Kapa Biosystems catalog # KK4824), and pooled for sequencing on an Illumina NextSeq 500 (2 × 75-bp). Reads were checked for quality using fastqc (11) and if necessary were trimmed using Trimmomatic (12) to trim regions with phred Q > 30 (13). High-quality reads were then aligned to reference genome hg19 using STAR (14). Gene counts were normalized by frequency per kilobase million (14). Differential expression of genes was determined using the limma (15) and DESeq2 (16) packages in the R environment (17), and multiple testing correction was performed using the FDR Benjamini–Hochberg method (18).

Principal components analysis (PCA) was used to visualize variation between samples based on expression of the top 2,000 most variably expressed genes within each cell line. One replicate sample from control-treated HCC-1428/LTED cells ("SH50") that was sequenced in a separate batch proved to be an outlier by PCA and hierarchical clustering; this sample was excluded from further analysis.

To determine significant gene expression pathway enrichment between time points, we conducted single-sample gene set enrichment analysis (ssGSEA) (19) for Hallmarks pathways in GenePattern (20) using default arguments. Gene set enrichment scores were normalized based on numbers of genes within each Hallmarks gene set. Normalized enrichment scores (NES) were compared between treatment groups by Bonferroni multiple comparison-adjusted posthoc test. RNA-seq data were deposited at NCBI Sequence Read Archive (SRA) under accession # PRJNA816494.

**Growth assays**

Parental cells were HD for 3 d prior to seeding. Cells were seeded in triplicate at 12,000 cells/well in 12-well plates, then treated as indicated for 4 wk. Cells were fixed and stained with 0.5% crystal violet in 20% methanol for 10 min. Excess dye was rinsed out with water. Plates were scanned, and area fraction of staining in each well was determined using ImageJ.

**Statistical analysis**

Cell growth data and IHC scores were analyzed by t-test (for 2-group experiments) or ANOVA followed by Bonferroni multiple comparison-adjusted post hoc testing between groups (for experiments with ≥3 groups). Pairwise comparisons of immunofluorescence scores used the Cramer-von Mises nonparametric test. Cell culture experiments were repeated on 3 independent occasions, and the data met the assumptions of all tests. Tumor growth data were analyzed using the following linear mixed model : $\text{Log}_{10}(\text{tumor volume}_{it}) = a_i + b * t + e_{it}$, where i represents the $i^{th}$ mouse, t represents time of tumor volume measurement, $a_i$ represents the mouse-specific log tumor volume at t = 0, b represents the rate of tumor volume growth, and $e_{it}$ represents deviation of measurements from the model over time (21,22). Mouse heterogeneity (the baseline tumor volume) is represented by variance of $a_i$, and $b*\log_e(10) * 100$ indicates tumor volume increase (%) per week. Treatment groups were compared using a z-test for slopes with standard error derived from the output of the function lme from the library nlme in R. Synergy was determined as described in ref. (7).

### Supplemental References Cited

1. Traphagen NA, Hosford SR, Jiang A, Marotti JD, Brauer BL, Demidenko E*, et al.* High estrogen receptor alpha activation confers resistance to estrogen deprivation and is required for therapeutic response to estrogen in breast cancer. Oncogene **2021**;40(19):3408-21.
2. Miller TW, Hennessy BT, Gonzalez-Angulo AM, Fox EM, Mills GB, Chen H*, et al.* Hyperactivation of phosphatidylinositol-3 kinase promotes escape from hormone dependence in estrogen receptor-positive human breast cancer. J Clin Invest **2010**;120(7):2406-13.
3. McQuin C, Goodman A, Chernyshev V, Kamentsky L, Cimini BA, Karhohs KW*, et al.* CellProfiler 3.0: Next-generation image processing for biology. PLoS Biol **2018**;16(7):e2005970.
4. DeRose YS, Gligorich KM, Wang G, Georgelas A, Bowman P, Courdy SJ*, et al.* Patient-derived models of human breast cancer: protocols for in vitro and in vivo applications in tumor biology and translational medicine. Curr Protoc Pharmacol **2013**;Chapter 14:Unit14 23.
5. Demidenko E. Mixed Models: Theory and Applications with R. Hoboken, NJ: Wiley; 2013.
6. Demidenko E. Advanced Statistics with Applications in R. Wiley (Hoboken, NJ). **2020**.
7. Demidenko E, Miller TW. Statistical determination of synergy based on Bliss definition of drugs independence. Plos One **2019**;14(11):e0224137.
8. Martin M. Cutadapt removes adapter sequences from high-throughput sequencing reads. 2011 **2011**;17(1):3.
9. Li W, Koster J, Xu H, Chen CH, Xiao T, Liu JS*, et al.* Quality control, modeling, and visualization of CRISPR screens with MAGeCK-VISPR. Genome biology **2015**;16:281.
10. Subramanian A, Tamayo P, Mootha VK, Mukherjee S, Ebert BL, Gillette MA*, et al.* Gene set enrichment analysis: a knowledge-based approach for interpreting genome-wide expression profiles. Proceedings of the National Academy of Sciences of the United States of America **2005**;102(43):15545-50.
11. Andrews S. FastQC: a quality control tool for high throughput sequence data. Available at: http://www.bioinformatics.babraham.ac.uk/projects/fastqc. **2010**.
12. Bolger AM, Lohse M, Usadel B. Trimmomatic: a flexible trimmer for Illumina sequence data. Bioinformatics **2014**;30(15):2114-20.
13. Ewing B, Hillier L, Wendl MC, Green P. Base-calling of automated sequencer traces using phred. I. Accuracy assessment. Genome research **1998**;8(3):175-85.
14. Dobin A, Davis CA, Schlesinger F, Drenkow J, Zaleski C, Jha S*, et al.* STAR: ultrafast universal RNA-seq aligner. Bioinformatics **2013**;29(1):15-21.
15. Ritchie ME, Phipson B, Wu D, Hu Y, Law CW, Shi W*, et al.* limma powers differential expression analyses for RNA-sequencing and microarray studies. Nucleic acids research **2015**;43(7):e47.
16. Love MI, Huber W, Anders S. Moderated estimation of fold change and dispersion for RNA-seq data with DESeq2. Genome biology **2014**;15(12):550.
17. R_Core_Team. R: A Language and Environment for Statistical Computing. R Foundation for Statistical Computing, Vienna. . **2017**.
18. Benjamini Y, Hochberg, Y. Controlling the false discovery rate – a practical and powerful approach to multiple testing. J R Stat Soc Series B Methodol **1995**;57:289-300.
19. Barbie DA, Tamayo P, Boehm JS, Kim SY, Moody SE, Dunn IF*, et al.* Systematic RNA interference reveals that oncogenic KRAS-driven cancers require TBK1. Nature **2009**;462(7269):108-12.
20. Reich M, Liefeld T, Gould J, Lerner J, Tamayo P, Mesirov JP. GenePattern 2.0. Nature genetics **2006**;38(5):500-1.
21. Demidenko E. Mixed models : theory and applications with R. Hoboken, New Jersey: Wiley; 2013. xxvii, 717 pages p.
22. Demidenko E. Advanced statistics with applications in R. Wiley series in probability and statistics. Hoboken, NJ: Wiley,; 2020. p 1 online resource.

**Supplemental Data**

**Fig. S1- E2 induces changes in overlapping sets of genes in HCC-1428 and HCC-1428/LTED cells.** (A) Differentially expressed genes (FDR<0.05) were identified within each cell line by comparison of transcriptomes from cells treated ± 1 nM x 24 h. Overlap analysis indicated that 851 genes were altered by E2 treatment in both cell lines. (B) Relative expression levels of these 851 genes in all samples were analyzed by unsupervised hierarchical clustering and are shown as a heatmap.

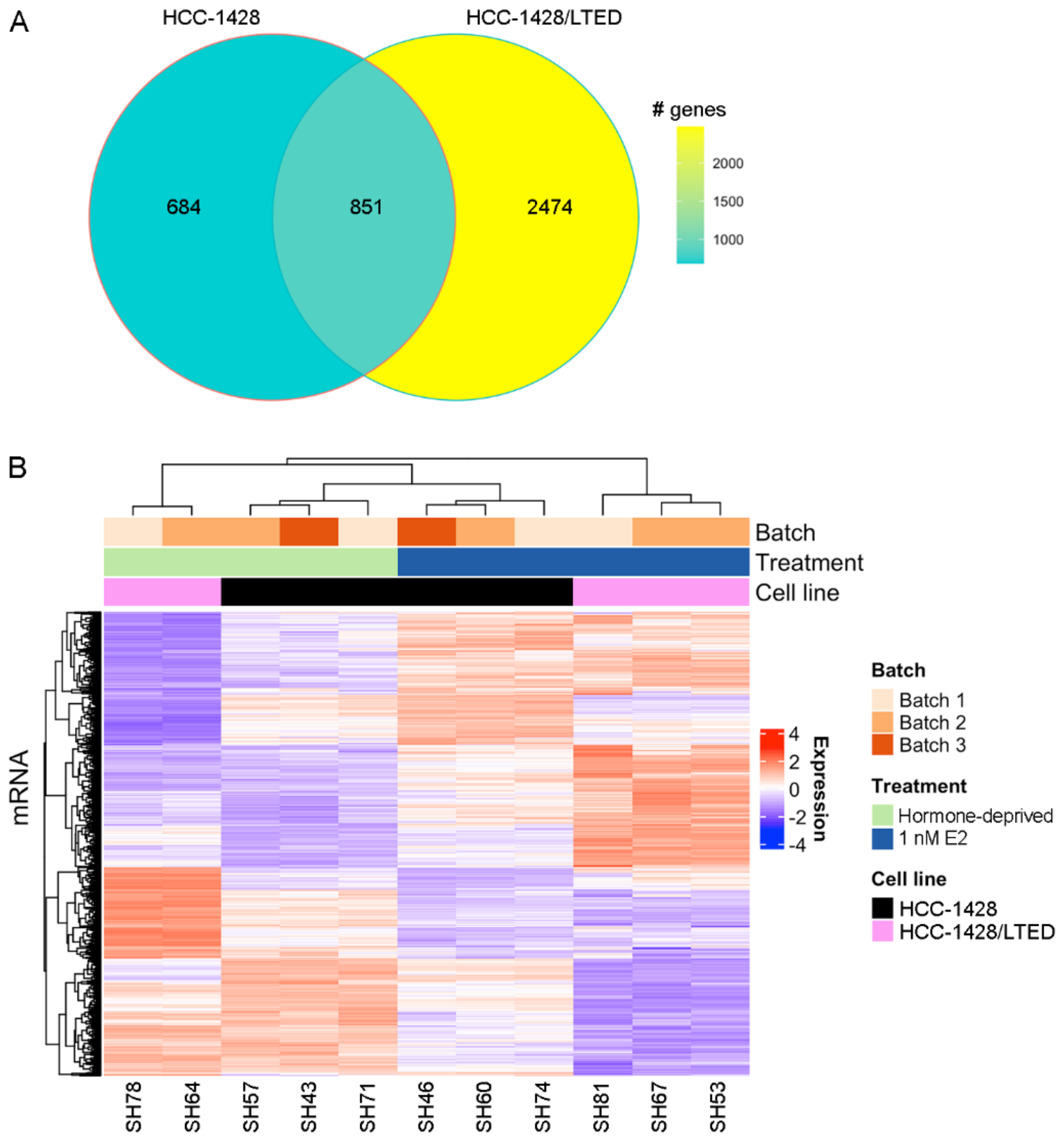

**Fig. S2- E2 induces DNA damage in LTED and ER-overexpressing cells.** In (A/B), cells were propagated in HD medium prior to seeding. In (C), T47D/pInd20-*ESR1* cells were pretreated with HD x 7 d, and then treated with HD + dox x 14 d prior to seeding. All cells were then treated as indicated x 24 h. Cells were fixed and stained for 53BP1 (green) and with DAPI (blue). Representative images are shown in (A). 53BP1 foci were counted in ≥100 cells per group. *p<0.05, ***p<0.0005, ****p<0.0001.

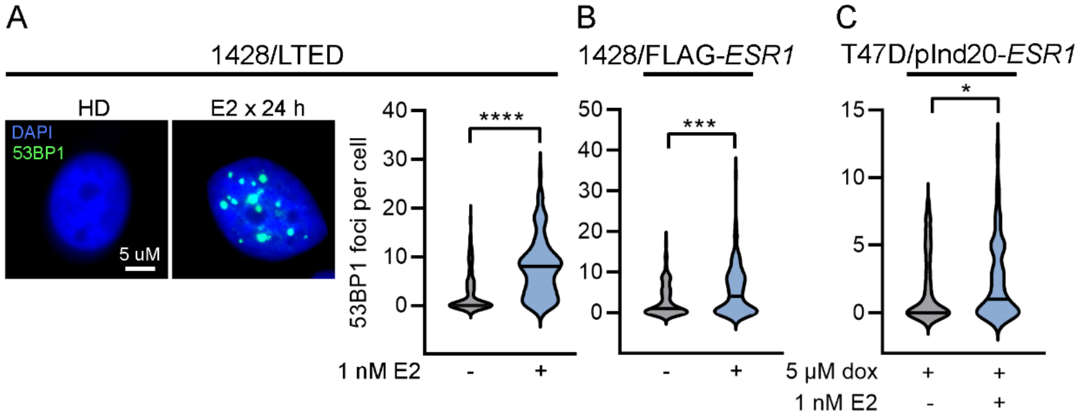

**Fig. S3- E2-induced DNA damage is dependent upon cell cycle progression.** (A/B) HCC-1428 and MDA-415/Luc cells were treated with HD x 3 d prior to seeding. All cell lines were then treated ± E2 for 21 h, and labeled with BrdU in culture medium for another 3 h prior to harvest. Floating and attached cells were harvested and stained for γH2AX, BrdU, and cleaved PARP, and with propidium iodide. Cells were analyzed by flow cytometry. Cleaved PARP-positive cells were considered apoptotic and excluded from analysis. Representative flow cytometry results are shown in (A). Proportions of cells with DNA breaks (γH2AX-positive) that were or were not in S-phase (i.e., did or did not incorporate BrdU) were plotted in (B). Data are shown as mean of triplicates + SD. Proportions of γH2AX+/BrdU+ cells were statistically compared between treatment groups as indicated with brackets. (C) T47D/pInd20-*ESR1* cells were pretreated with HD x 7 d, and then treated with HD + dox x 14 d prior to seeding. All cell lines were then treated ±E2 ± abemaciclib as indicated x 24 h, fixed, and stained for γH2AX and with DAPI. γH2AX foci were counted in ≥100 nuclei per group. *p<0.05, **p<0.005, ****p<0.0001.

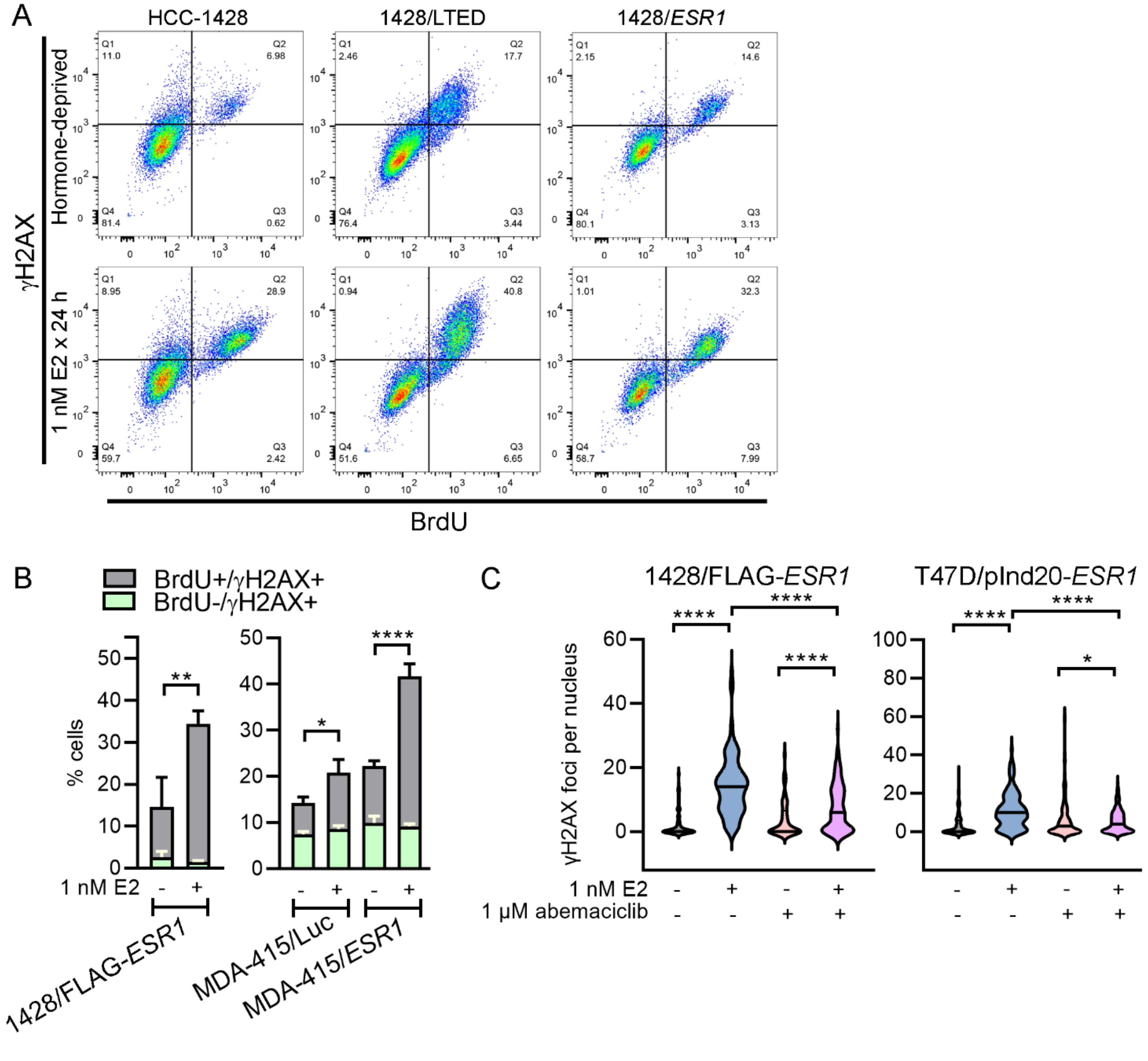

**Fig. S4- ER overexpression converts E2 from a growth promoter to a growth suppressor.** (A) Lysates from hormone-deprived HCC-1428 (HD x 3 d), HCC-1428/LTED, HCC-1428/vec (HD x 3 d), HCC-1428/FLAG-ESR1, MDA-MB-415/vec, MDA-MB-415/FLAG-*ESR1*, and T47D/pInd20-ESR1 cells (HD x 3 d, then treated ± dox x 2 d) were analyzed by immunoblot. (B) HCC-1428/vec and MDA-MB-415/vec cells (HD x 3 d), and HCC-1428/FLAG-*ESR1* and MDA-MB-415/FLAG-*ESR1* cells (propagated in HD) were seeded and then treated ± E2 for 4 wk. T47D/pInd20-ESR1 cells (HD x 3 d) were treated ± dox for 2 wk, then reseeded and treated ± dox ± E2 for 2 wk. Relative cell growth was measured. Data are shown as mean of triplicates ± SD. (C) HCC-1428/LTED cell lines expressing dox-inducible shRNA targeting *ESR1* (two independent constructs) or non-silencing control were treated ± dox for 2 d, and lysates were analyzed by immunoblot. *p<0.05, **p<0.005, ***p<0.0005.

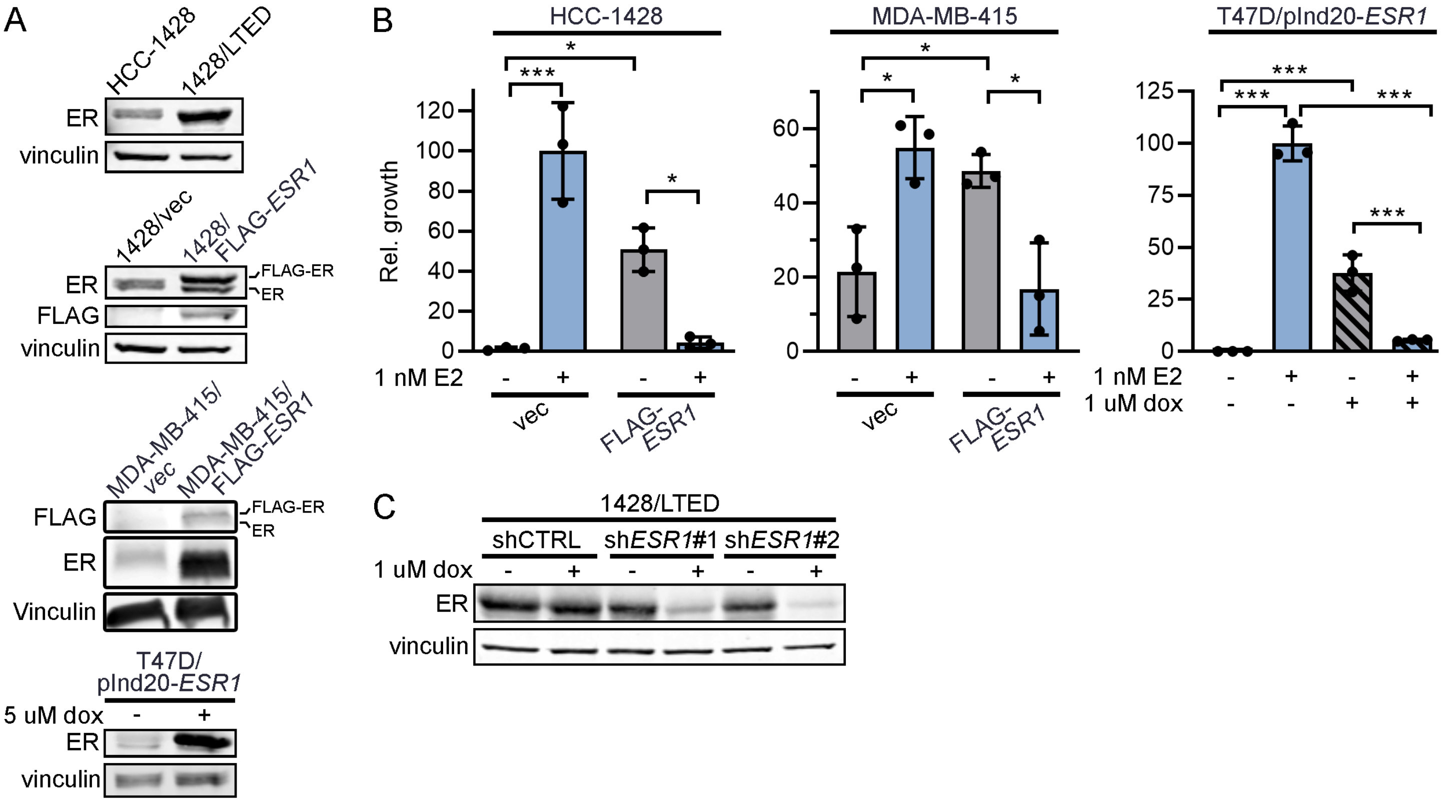

**Fig. S5- ER-negative BT-20 cells do not exhibit increased DNA damage upon E2 treatment.** BT-20 cells were HD x 3 d, then treated ± 1 nM E2 x 24 h. Cells were fixed and stained for γH2AX (green) and with DAPI (blue). Representative images are shown. γH2AX foci were counted in ≥100 cells per group. n.s. = not significant.

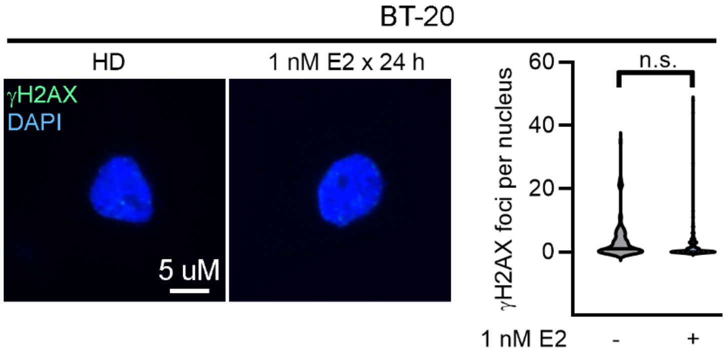

**Fig. S6- Histology of baseline human tumors.** Representative images of H&E, ER, and PR IHC staining of human tumor specimens (analyzed in Fig. 3A) obtained prior to E2 treatment are shown.

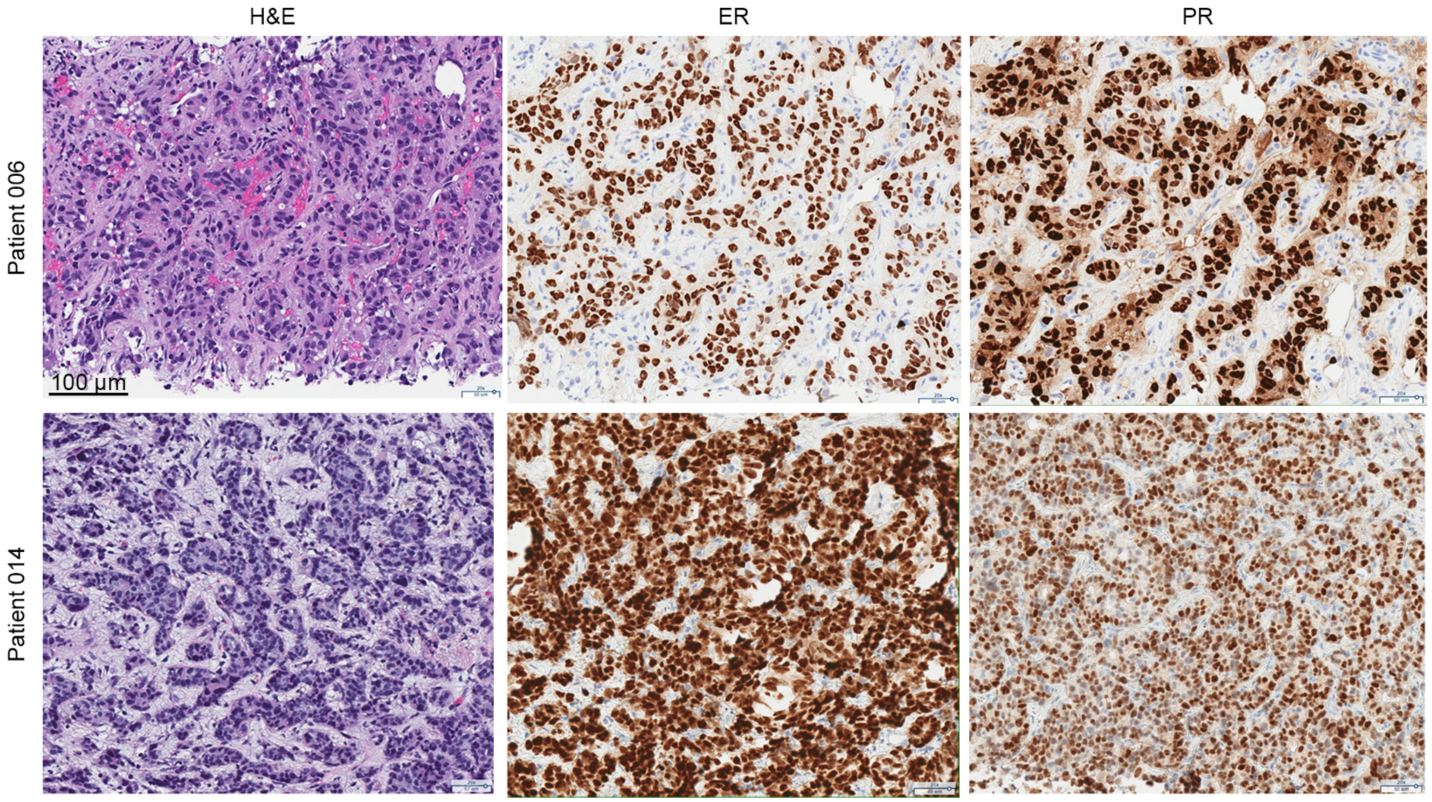

**Fig. S7- Single-channel fluorescence images of tumors from patients treated with E2.** Exposure-matched paired images are shown. Merged images are shown in Fig. 3A.

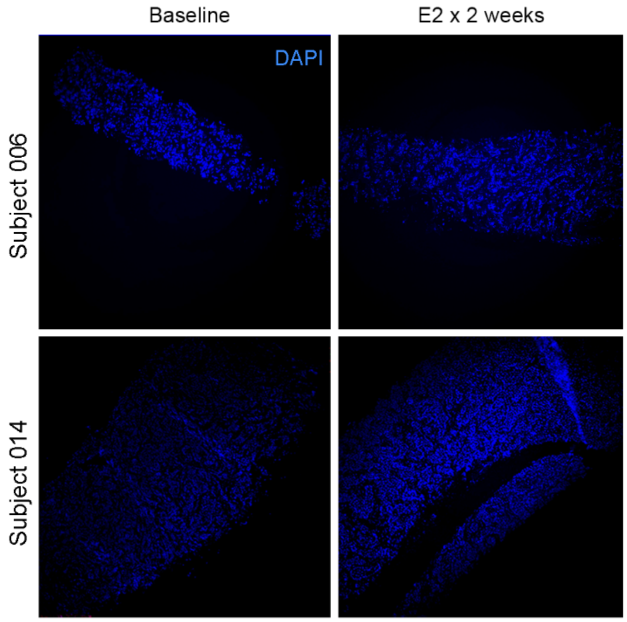


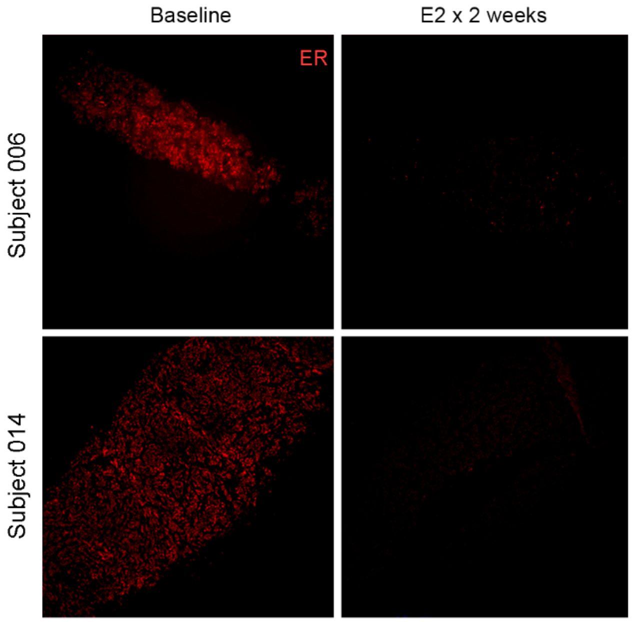


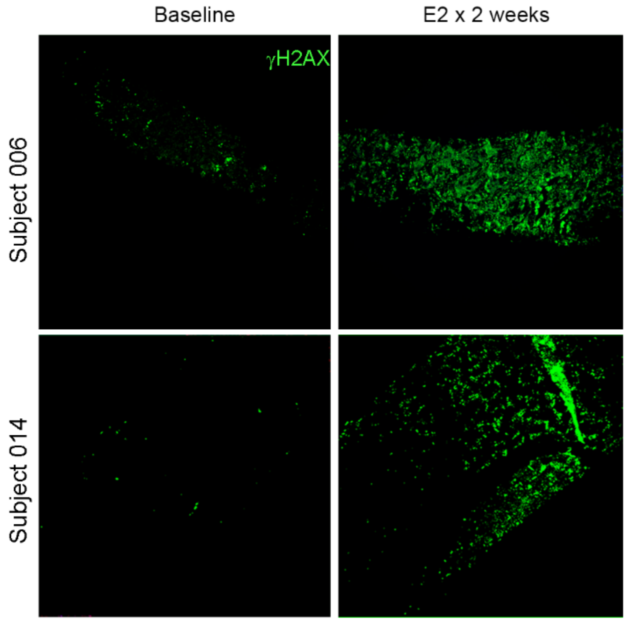

**Fig. S8- Individual tumor volumes from mice treated ± E2.** Ovx mice bearing tumors ~200 $mm^3$ were randomized to treatment ± E2 by s.c. pellet. Tumor volumes were serially measured. Each line represents one mouse. Summary data are shown in Fig. 3B.

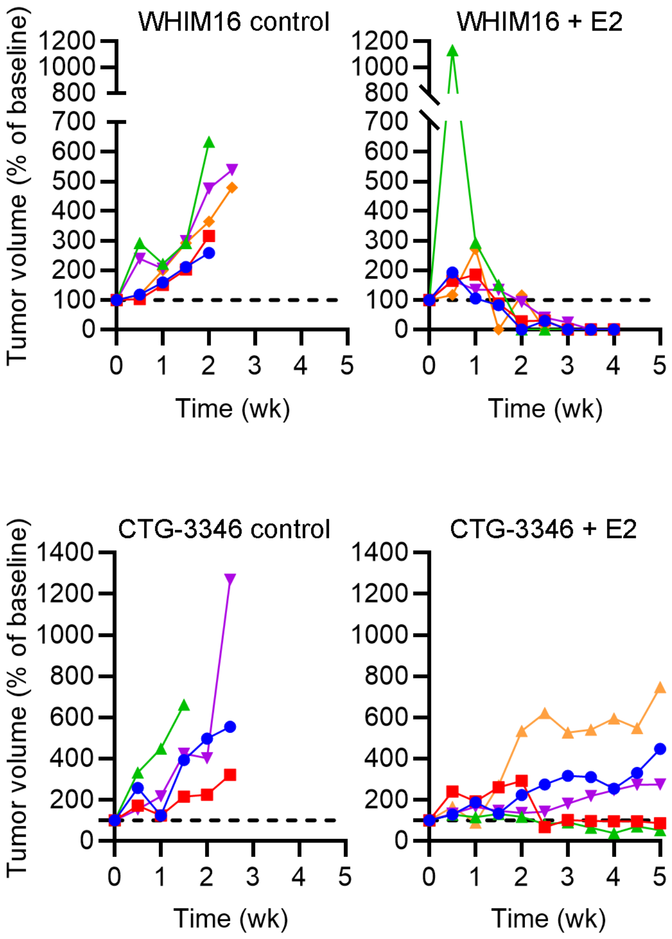

**Fig. S9- Characterization of PDX model CTG-3346.** (A) Treatment history of the patient that provided tumor tissue for PDX. (B) Representative H&E, ER, and PR IHC staining of patient's tumor before and after (P3) growth in mice. (C) Tumors were harvested from ovx NSG mice. Tumor lysates were analyzed by immunoblot. WHIM16 was included as a positive control for Rb expression. (D) Ovx mice bearing CTG-3346 tumors ~200 $mm^3$ were randomized to treatment ± E2 (s.c. pellet) for 24 h. RNA was isolated from tumors, and gene expression was measured by RT-qPCR. *TFF1* levels were normalized to *ACTB* levels. Horizontal bars indicate median values. *p<0.05.

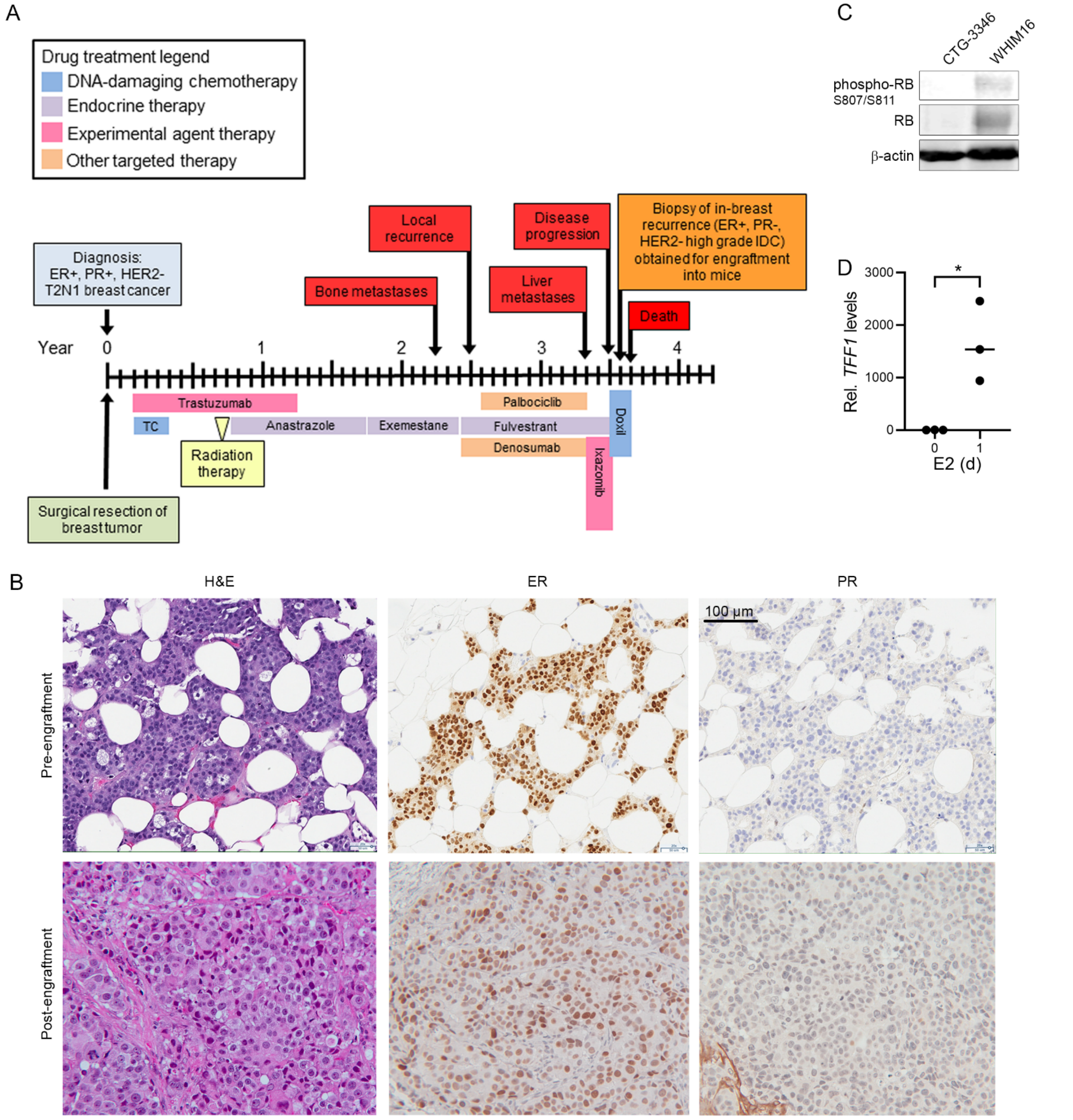

**Fig. S10- Representative IHC images from WHIM16 and CTG-3346 tumors.** Ovx mice bearing tumors ~200 $mm^3$ were randomized to treatment ± E2 by s.c. pellet. Tumors were harvested 24 h later and FFPE. Sections were analyzed by IHC. Quantitative results are shown in Fig. 3C.

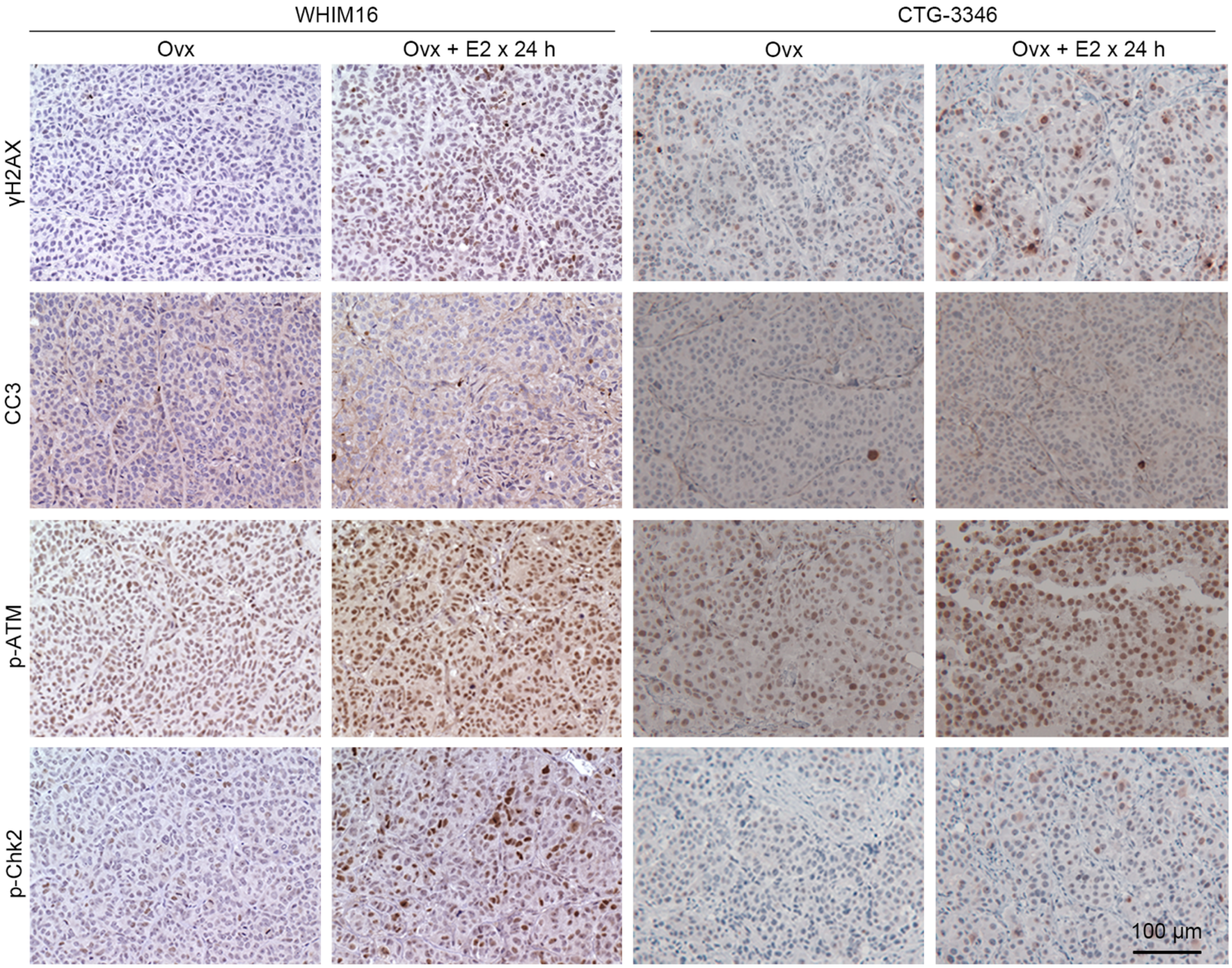

**Fig. S11- E2 stimulation induces DNA replication stress.** T47D/pInd20-*ESR1* cells were pretreated with HD x 7 d, and then treated with HD + dox x 14 d prior to seeding. All cells were then treated as indicated x 24 h. Cells were fixed and stained for PCNA and with DAPI. PCNA foci were counted in ≥100 cells per group. ****p<0.001.

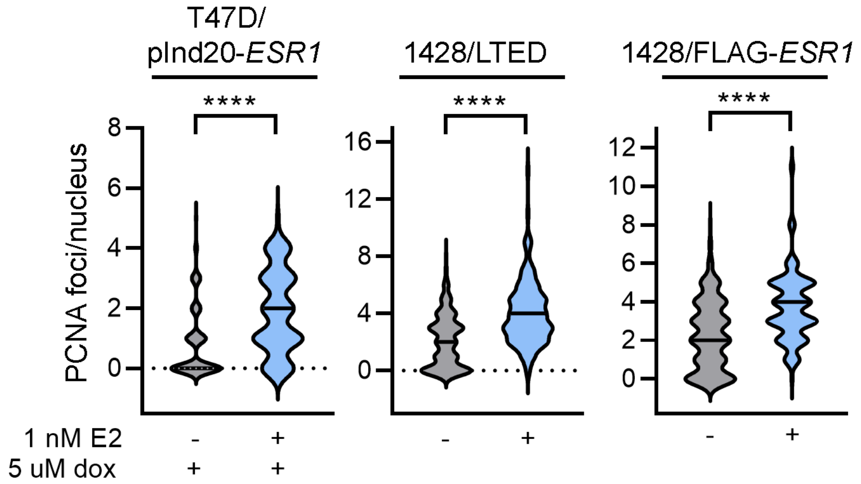

**Fig. S12- ER overexpression promotes E2-induced R-loop formation.** (A/B) T47D/pInd20-*ESR1* cells were pretreated with HD x 7 d, and then treated with HD + dox x 14 d prior to seeding. Both cell lines were then treated as indicated x 24 h, fixed, and stained for DNA/RNA hybrids (S9.6 antibody) and with DAPI. S9.6 fluorescence intensity was quantified in ≥100 nuclei per group. (C) Cells were transiently transfected with plasmids encoding RNase H1 or vector control. Two days later, cells were treated ± 1 nM E2 x 24 h, then fixed, stained, and analyzed as in (B). Representative images are shown. *p<0.05, **p<0.005, ***p<0.0005, ****p<0.0001 compared to control unless otherwise indicated.

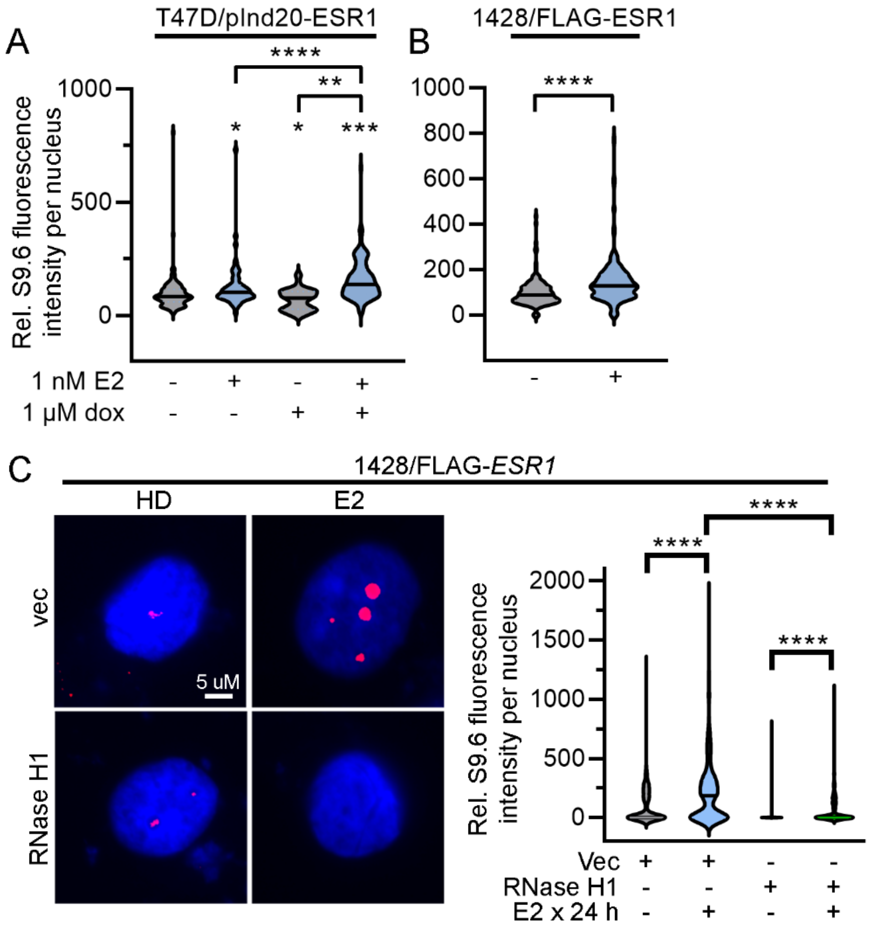

**Fig. S13- E2 and olaparib combine to increase R-loop formation.** Cells were treated ± olaparib for 2 d, then treated ± olaparib ± E2 x 24 h. Cells were fixed and stained for DNA:RNA hybrids (S9.6 antibody) and with DAPI. S9.6 fluorescence intensity was quantified in ≥100 nuclei/group. *p<0.05, ****p<0.0001.

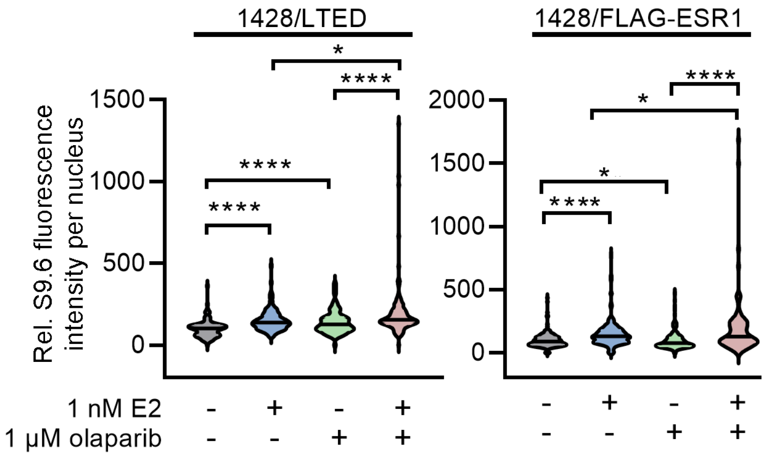

**Fig. S14- Individual growth curves of tumors from mice treated ± E2 and/or olaparib.** Ovx NSG mice bearing tumors ~200 $mm^3$ were randomized to treatment with vehicle, olaparib (50 mg/kg/d i.p.), E2 (1 mg s.c. pellet, replaced every 30 d), or the combination of E2 and olaparib. E2 was administered continuously. Olaparib was administered daily for 28 d, then stopped. Tumor volumes were serially measured. Each line represents one mouse. Summary data are shown in Fig. 6A/B.

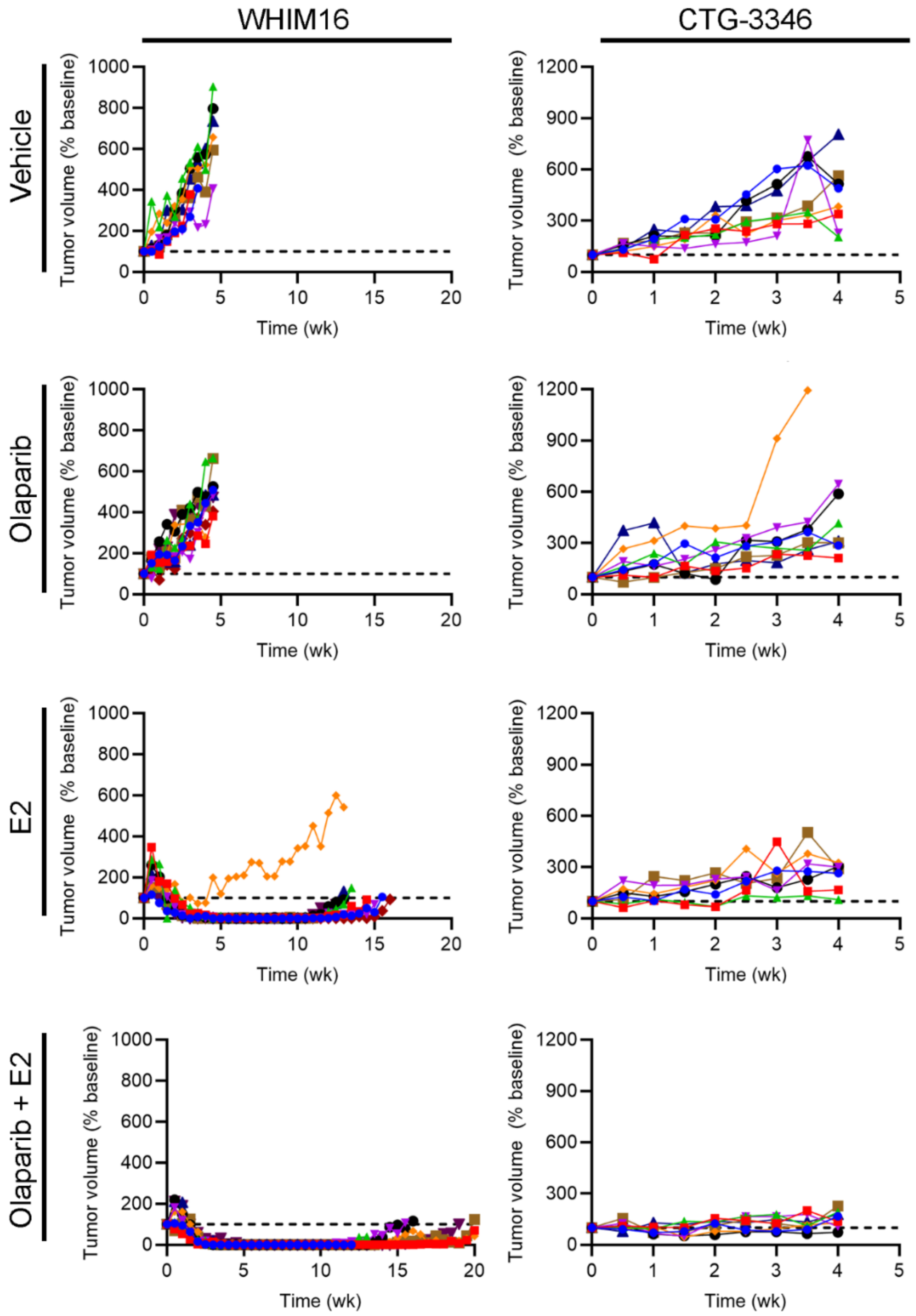

**Fig. S15- Individual growth curves of WHIM16 tumors from mice treated with a second cycle of E2 ± olaparib.** Mice bearing WHIM16 tumors that recurred on E2 monotherapy, and then resumed growth during subsequent estrogen deprivation (in Fig. 6D), were randomized to treatment with E2 (1 mg s.c. pellet) ± olaparib (50 mg/kg/d i.p.). Tumor volumes were serially measured. Each line represents one mouse. Summary data are shown in Fig. 6E.

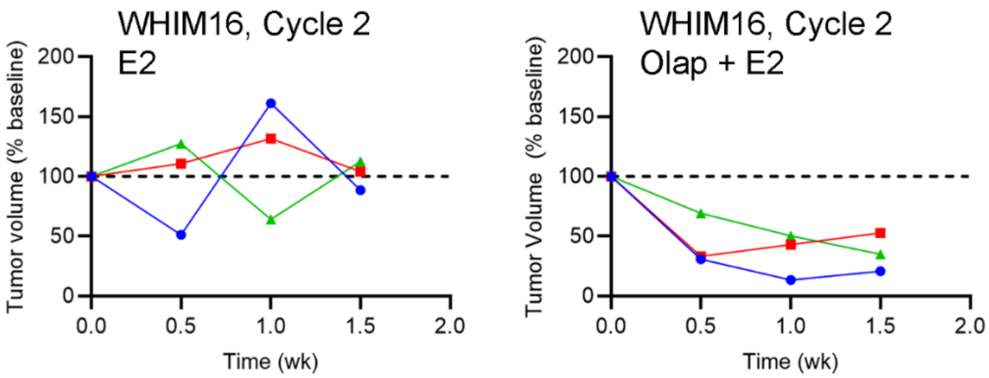
